# Alternative splicing of SLAMF6 in human T cells creates a co-stimulatory isoform that counteracts the inhibitory effect of the full-length receptor

**DOI:** 10.1101/2020.08.21.262238

**Authors:** Emma Hajaj, Elad Zisman, Shay Tzaban, Sharon Merims, Jonathan Cohen, Shiri Klein, Shoshana Frankenburg, Moshe Sade-Feldman, Yuval Tabach, Keren Yizhak, Ami Navon, Polina Stepensky, Nir Hacohen, Tamar Peretz, André Veillette, Rotem Karni, Galit Eisenberg, Michal Lotem

**Affiliations:** Sharett Institute of Oncology, Hadassah Hebrew University Hospital, Jerusalem, Israel; Wohl institute for translational medicine, Hadassah Medical Organization, Jerusalem, Israel; Lautenberg center for immunology and cancer research, Faculty of Medicine, Hebrew University, Jerusalem, Israel; Department of Cell Biology and Cancer Science, Rappaport Faculty of Medicine, Technion–Israel Institute of Technology, Haifa, Israel; Broad Institute of MIT and Harvard, Cambridge, MA 02142, USA; Department of Medicine, Center for Cancer Research, Massachusetts General Hospital, Boston, MA 02114, USA; Department of Medicine, Harvard Medical School, Boston, MA 02115, USA; Department of Developmental Biology and Cancer Research, Institute for Medical Research Israel-Canada, Hebrew University of Jerusalem; Department of biological regulation, The faculty of biology, Weizmann institute of science, Rechovot, Israel; Department of bone marrow transplantation, Hadassah Medical Organization, Jerusalem, Israel; IRCM, Montreal Clinical Research Institute, Montreal, Canada; Department of Biochemistry and Molecular Biology.The Institute for Medical Research Israel-Canada, Hebrew University-Hadassah Medical School, Jerusalem, Israel

## Abstract

SLAMF6 is a homotypic receptor of the Ig-superfamily associated with progenitor exhausted T cells. In humans, SLAMF6 has three splice isoforms involving its V-domain. While the canonical 8-exon receptor inhibits T cell activation through SAP recruitment, the short isoform SLAMF6^Δ17-65^ has a strong agonistic effect. The costimulatory action depends on protein phosphatase SHP-1 and leads to a cytotoxic molecular profile governed by transcription factors Tbet, Runx3, and Tcf7. In T cells from individual patients treated with immune checkpoint blockade, a shift was noted towards SLAMF6^Δ17-65^. Splice-switching antisense oligonucleotides designed to target the SLAMF6 splice junction, enhanced SLAMF6^Δ17-65^ in human tumor-infiltrating lymphocytes and improved their capacity to inhibit human melanoma in mice. The possible emergence of two opposing isoforms from the SLAMF6 gene may represent an immune-modulatory mechanism that can be exploited for cancer immunotherapy.

## Introduction

The SLAM family of immune receptors (SFR) comprises nine genes located on chromosome 1 and is considered part of the CD2 cluster of the immunoglobulin superfamily (IgSF)^1,2^. Most members of this group are self-binding receptors engaged in homotypic interactions; they cluster at the immune synapse^3^ and serve as collectivity sensors^4^. SFRs are varyingly expressed on cells of hematopoietic progeny, and often more than one SFR is present on a cell subset. SFRs participate in lymphocyte and NKT development, and B -T cell interactions, but most notably, they co-modulate B, T, and NK cell activation^1,4,5,6,7^. It is characteristic of SFRs that at least one unique immunoreceptor tyrosine-based switch motif (ITSM) appears in their cytoplasmic portion^8^. The amino acid sequence motif TxYxxV/I can recruit the SH2-homology domain-containing adaptor proteins SAP and EAT2^9,10^, or protein phosphatases SHP-1, 2, and SHIP^11^. The capacity of SFRs to acquire two types of adaptors, one that recruits kinases or one that de-phosphorylates signaling molecules, has supported the idea of a dual function^3^. The duality concept was corroborated by studies of inborn X-linked lymphoproliferative disease, in which a mutation in SAP leads to severe immune dysregulation^12^.

In contrast to the paradigm of a bidirectional modulatory role of SFRs, we recently reported that genetic deletion of SLAMF6 in melanoma-specific murine CD8 T cells significantly enhances their functional capacity^13^. We showed that the SLAMF6 KO-melanoma antigen-specific T cells demonstrated a global improvement of effector function, including cytokine release, cytotoxicity, and post-transfer persistence. Interestingly, activated SLAMF6 KO cells expressed low to null levels of SAP protein, while phosphatase SHP-1 was intact. We concluded that SLAMF6 is an obligatory negative checkpoint receptor and suggested that the acquisition of SAP is required for the inhibitory effect. In contrast, there was no indication that SHP-1 is necessary for SLAMF6-mediated co-inhibition

The main limit to extrapolating from the functional data generated with murine SLAMF6 (also known as Ly108) to the human receptor, relates to a different splicing pattern. The two murine Ly108 splice variants occur in the cytoplasmic region. Ly108-1 contains one tyrosine-based motif, and Ly108-2 bears two motifs^14^. Intriguingly, Ly108-1 is associated with a propensity to develop autoimmune systemic lupus erythematosus (SLE)^15^. In contrast to mouse SLAMF6, splicing of the human SLAMF6 molecule occurs in the extracellular part of the receptor: the canonical protein includes eight exons with Ig-like V and C2 domains; the isoform SLAMF6^Δ17-65^ lacks part of exon2, and SLAMF6^Δ18-128^ is an isoform in which the entire exon2 is skipped (NCBI Reference Sequences: NM_001184714.2, NM_001184715.2, NM_001184716.2).

The fact that the human SLAMF6 ectodomain, the critical part for signal initiation, appears in molecular configurations that do not exist in mice, prompted us to study human SLAMF6 splice variants. Surprisingly, we found that while the canonical SLAMF6 acts as a co-inhibitory receptor, SLAMF6^Δ17-65^ functions as a strong co-stimulatory agonist. By producing T cells that do not express the canonical SLAMF6, we were able to study in-depth the molecular events promoted by the shorter isoform. On the basis of that study, we developed a method based on antisense oligonucleotides (ASOs) that was implemented for adoptive cell therapy of melanoma.

## Results

### Identification of SLAMF6 splice isoforms on T cell subsets

The human SLAMF6 gene was cloned by the Moretta group in 2001 and mapped in the SLAM gene cluster on Chr 1q22 by Terhorst, in 2002^16,17^. The gene encodes a 331 amino acid sequence with four splice isoforms recorded on RefSeq^18^. Isoform 1 is the canonical sequence composed of an immunoglobulin core with variable and constant domains (NCBI Reference Sequence: NM_001184714.2). Isoform 3 includes an alternative acceptor site, which consists of a 3’ alternative splicing of exon2, lacking amino acids 17-65 of the variable region (NM_001184715.2), labeled here SLAMF6^Δ17-65^. Isoform 4 is the result of exon2 removal, which leads to a protein missing the whole variable domain, labeled here SLAMF6^ΔExon2^ (NM_001184716.2, Figure 1a, b). The transmembrane and intracellular domains of these three isoforms are identical. Another isoform of SLAMF6, which is inseparable from isoform 1, has one alanine missing in the intracellular domain but intact immune tyrosine domains [two immune tyrosine switch motifs (ITSMs) and one immune tyrosine inhibitory domain (ITIM)].

**Figure 1:**
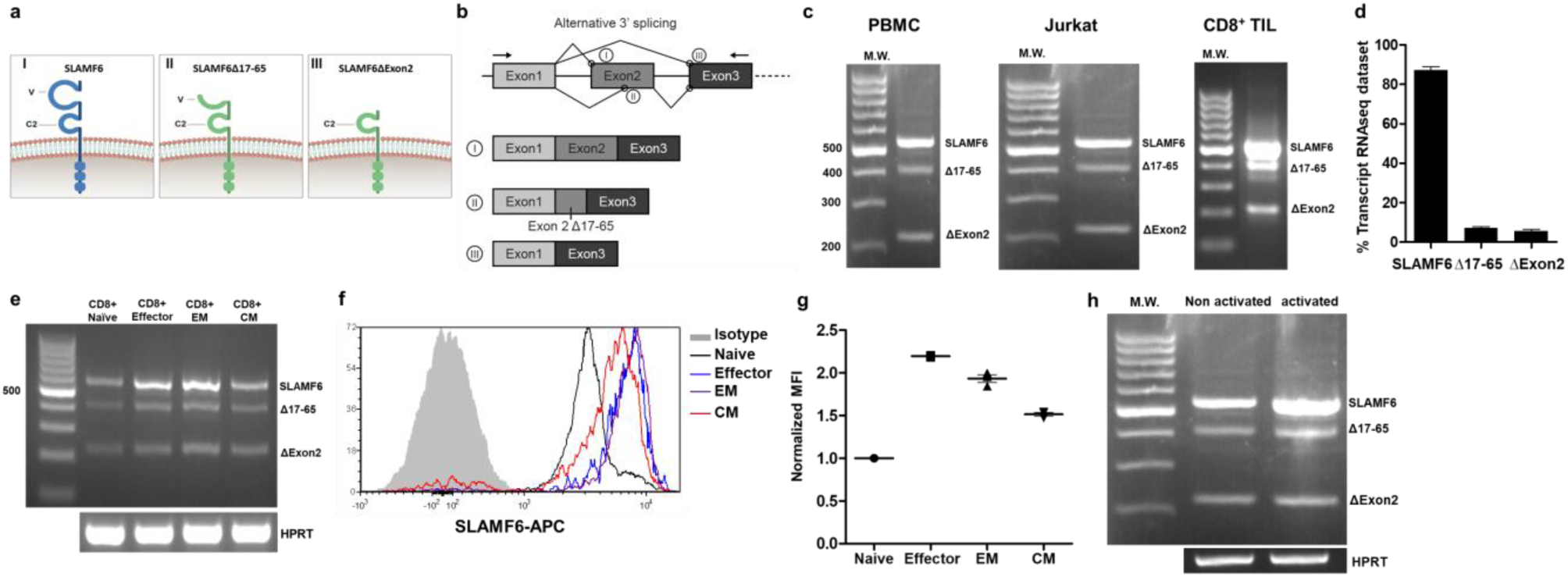
Identification of SLAMF6 splice isoforms in T cell subsets. **a, b**, SLAMF6 isoforms: (I) Canonical full-length isoform. (II) SLAMF6^Δ17-65^ 3’ splice isoform; (III), SLAM6^Δexon2^. **a**, Schematic diagram of each isoform on the membrane, **b**, Schematic diagram of exon2 alternative splicing. **c**, Expression of *SLAMF6* isoforms in peripheral blood mononuclear cells (PBMC), Jurkat cells, and tumor-infiltrating lymphocytes (TIL). **d**, Expression of *SLAMF6* splice isoforms in T cells from healthy donors (Accession: PRJNA482654). **e - g**, SLAMF6 isoform in naϊve, effector, EM, and CM CD8 T cell subsets. **e**, RNA expression. **f**, Flow cytometry of SLAMF6. **g**, Median fluorescence intensity (MFI) of SLAMF6. EM, effector memory. CM, central memory. **h**, RNA expression of *SLAMF6* isoforms in 48-h activated and non-activated Jurkat cells.

Using primers that complement sequences upstream and downstream of exon2 splice sites, we identified 3 bands, matching isoforms 1, 3, and 4, as referenced in RefSeq, by their correct size, in RNA extracted from PBMCs (peripheral blood mononuclear cells), Jurkat cells, and CD8+ tumor-infiltrating lymphocytes (TILs) (Figure 1c). We sequenced the bands to confirm their identity.

To confirm the existence of these isoforms, we analyzed a public RNA sequencing dataset (Accession: PRJNA482654)^19^. Using the RSEM software package for quantifying isoform abundance, we identified all *SLAMF6* RNA isoforms (Figure 1d). We failed to identify exon2 splicing in the murine analog of SLAMF6, *Ly108*, nor did we find data on murine isoforms in *Ly108* extracellular domain in genome browsers (Supplementary figure 1a). We next asked whether the splicing of *SLAMF6* varies in T cells during differentiation and activation. The *SLAMF6* canonical isoform, isoform 1, was upregulated in effector cells and effector memory cells compared to naϊve cells or to central memory (CM) cells (Figure 1e, Supplementary figure 1b), but we couldn’t detect a difference in the splicing ratio. Flow cytometry confirmed that SLAMF6 protein increased in parallel (Figure 1f,g). To test for isoform expression during activation, we used RNA from 48h-activated Jurkat cells (Figure 1h) and identified a general rise in *SLAMF6* transcripts. In conclusion, we found that the *SLAMF6* mRNA increased in activated, effector T cells with a corespondent amplification of all its isoforms.

### Trans-binding of full-length SLAMF6 on activated T cells inhibits their function while SLAMF6^Δ17-65^ acts as a co-stimulator

SLAMF6, a self-binding receptor, is expressed on hematopoietic cell lineages^1,2,20,21^ but not on non-hematopoietic cells. To evaluate the self-binding properties of SLAMF6 isoforms *in trans*, we took advantage of the receptor’s absence in somatic cells. We ectopically expressed isoform 1 (full length), isoform 3 (Δ17-65), and isoform 4 (ΔExon2) in a melanoma cell line (Supplementary figure 2a,b). We co-cultured the SLAMF6-expressing 526mel-SLAMF6 (full-length), 526*mel*-Δ17-65 and 526*mel*-ΔExon2 with cognate CD8 TILs. Compared to the original melanoma cells, melanoma cells expressing canonical SLAMF6 secreted significantly less IFN-γ (p<0.001), while TILs co-cultured with melanoma cells expressing SLAMF6^Δ17-65^ secreted higher levels of IFN-γ (p<0.05). Aberrant expression of SLAMF6^ΔExon2^, which lacks the binding interface of the receptor, had no effect on IFN-γ secretion in the co-culture assays of melanoma and TILs (Figure 2a-c).

**Figure 2:**
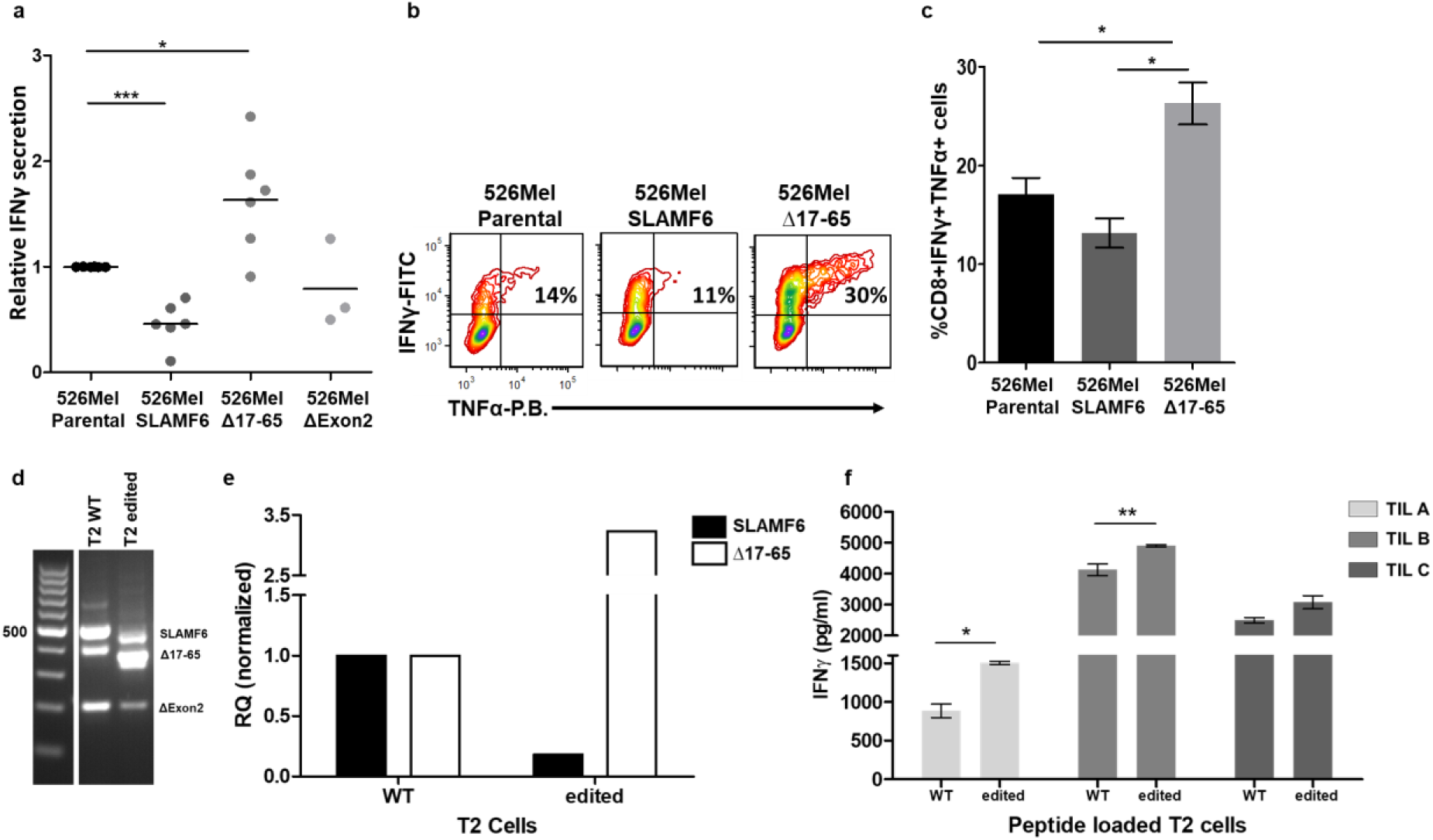
Trans-binding of full-length SLAMF6 on activated T cells inhibits their function while SLAMF6^Δ17-65^ acts as a co-stimulator. **a**, IFN-γ secretion by three cognate TILs incubated overnight with 526*mel* melanoma cell lines aberrantly expressing SLAMF6, SLAMF6^Δ17-65^ or SLAM6^Δexon2^. **b-c**, Flow cytometry for IFN-γ and TNF-α secretion by TIL209 melanoma-cognate CD8 T cells following 6 hours incubation with 526*mel* melanoma cell lines aberrantly expressing SLAMF6 isoforms. **d-f**, Stimulation of cognate CD8 TILs by antigen-presenting SLAMF6-edited T2 cells. **d-e**, SLAMF6-RNA extracted from CRISPR-Cas9 edited T2 cells. **f**, IFN-γ secretion by 3 cognate TILs following co-culture with SLAMF6-modified T2 or WT T2 cells pulsed with melanoma-specific peptides. One-way ANOVA. *, P < 0.05, ***, P < 0.001.

To support this observation, we established another model for *trans* activation using TAP-deficient T2 cells, which naturally express SLAMF6. Using CRISPR-Cas9*-*directed mutations of the SLAMF6 exon2 splicing site, we generated a single*-*cell clone with a higher expression of SLAMF6^Δ17-65^ compared to the canonical isoform (Figure 2d,e). When pulsed with a cognate melanoma peptide identified by three TILs, TILs co-cultures with the T2 cells expressing higher levels of SLAMF6^Δ17-65^ secreted more IFN-γ than the TILs co-cultured WT T2 cells. (Figure 2f).

Terhorst and our group have previously shown that canonical SLAMF6 serves as an inducer of T cell exhaustion and acts as an inhibitory checkpoint for T cells^13,22,23^. The results presented here with the full-length SLAMF6 support this role. Surprisingly, SLAMF6^Δ17-65^ seemed to be a counterpoise to full-length SLAMF6, acting as a co-stimulator *in trans*.

### Self-binding and cross-binding of SLAMF6 isoform ectodomains

After showing the effect of SLAMF6 isoforms, acting as ligands of their corresponding receptors on T cells, we asked if the isoforms pair together and if SLAMF6^Δ17-65^ can cross-bind the canonical receptor. To test the possibility that the canonical SLAMF6 associates with the extracellular domains of both the full-length molecule and of SLAMF6^Δ17-65^, we used a binding competition assay with HEK293 cells ectopically expressing one isoform of SLAMF6 (Figure 3a,b), and incubated the cells in the presence of the soluble ectodomain of SLAMF6 (seSLAMF6) in increasing concentrations to compete with a blocking antibody of SLAMF6, detectable by flow cytometry. Also, a quantitative ELISA was performed. Soluble ectodomains of SLAMF6 and SLAMF6^Δ17-65^ served either as a plate-bound bait or as a secondary biotinylated protein detectable with HRP-conjugated streptavidin (Figure 3c). The self-binding capacity of seSLAMF6^Δ17-65^ was the strongest, while the self-binding capacity of seSLAMF6 was the weakest. The pairing of seSLAMF6 and seSLAMF6^Δ17-65^ was stronger than SLAMF6 homotypic binding.

**Figure 3:**
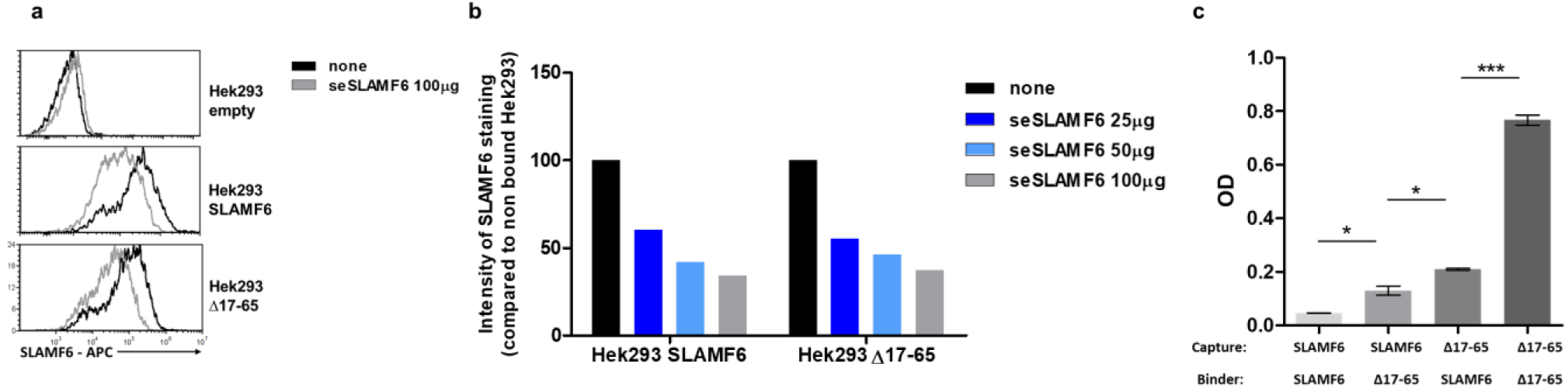
Self-binding and cross-binding of SLAMF6 isoform ectodomains. **a, b**, HEK293 cells were transfected with constructs of SLAMF6 isoforms or empty construct. The transfected cells were cultured with the soluble ectodomain of canonical SLAMF6 in the specified concentrations, washed, and stained for flow analysis of SLAMF6 expression using an anti-SLAMF6 antibody targeting a sequence in the C’ domain of the protein. **a**, Histograms showing SLAMF6 expression in each cell line after addition of 100 μg soluble SLAMF6, or no addition. **b**, Summary showing the MFI of SLAMF6 staining for every condition. **c**, ELISA showing the self-binding and cross-binding of soluble ectodomains of canonical SLAMF6 and SLAMF6^Δ17-65^. One-way ANOVA. *, P < 0.05, ***, P < 0.001.

We concluded from these experiments that the ectodomains of SLAMF6 and SLAMF6^Δ17-65^ cross-react, with the latter displaying a more robust binding capacity.

### Gene editing to delete canonical SLAMF6 while preserving SLAMF6^Δ17-65^/SLAMF6^Δexon2^ improves T cell function

The opposing co-modulatory effects of the canonical SLAMF6 and SLAMF6^Δ17-65^ when expressed on target cells as ligands, the first inhibiting T cells, and the latter co-stimulating, raised the question of the contribution of each isoform when acting as a receptor. To address this issue, we generated T cells expressing only the short isoforms of SLAMF6 (isoforms 3 and 4) using CRISPR-Cas9 targeting exon2 in the part that is skipped-out in Δ17-65 and Δexon2 (Figure 4a). Thus, protein translated from mRNA that includes mutated exon2 will not be viable, while transcripts in which the mutated segment is skipped by alternative splicing will be appropriately translated. For stable transfectants, we used a single-cell colony generated from the Jurkat T cell line following CRISPR-Cas 9 editing, in which a frameshift mutation leading to a stop-codon occurred. The cells were named Jurkat-SLAMF6^Δ17-65^ cells for the functional significance of this variant. Of note, isoform 4 (SLAMF6^Δexon2^), which showed no effect *in trans*, was not deleted, as there is no possibility to remove this isoform without eradicating isoform 3 (SLAMF6^Δ17-65^).

**Figure 4:**
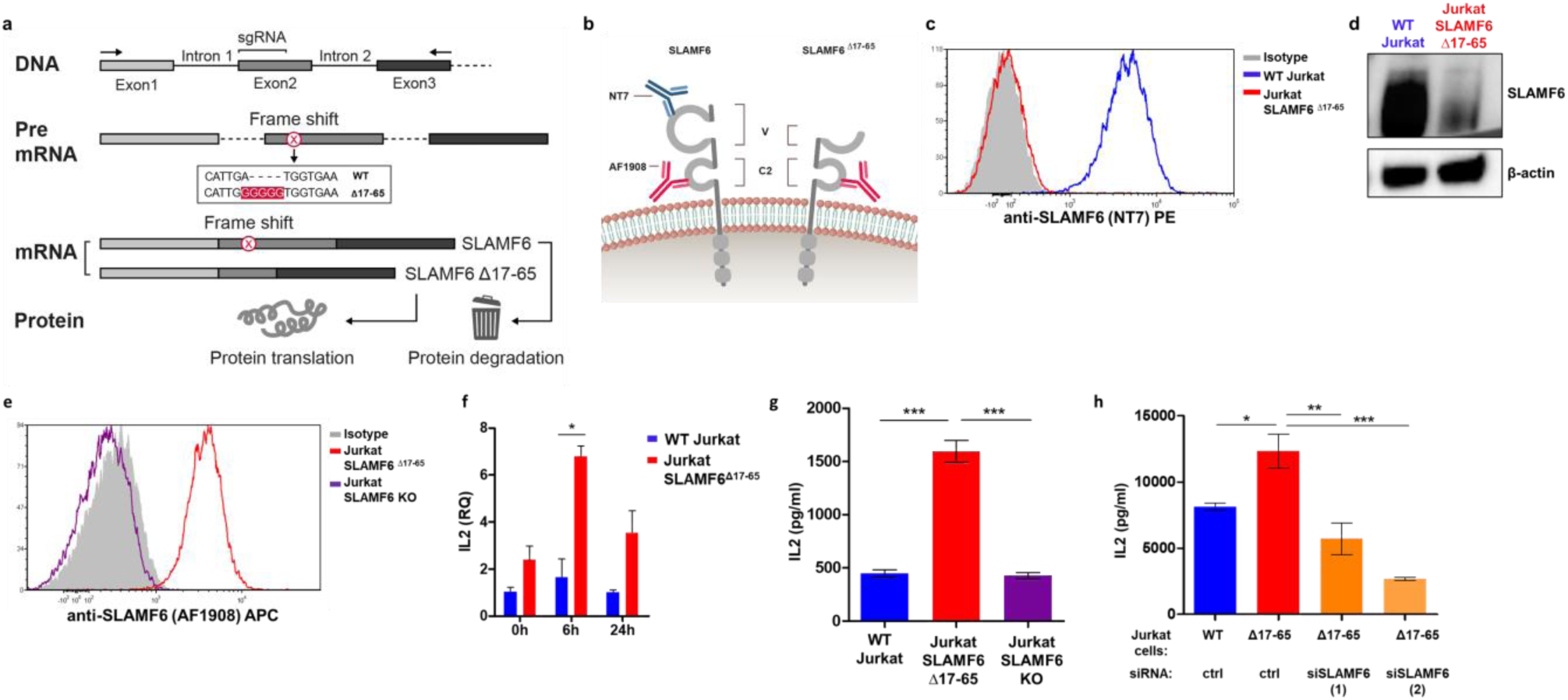
Gene editing to delete canonical SLAMF6 while preserving SLAMF6^Δ17-65^/ SLAMF6^ΔExon2^ improves T cell function. **a**, sgRNA targeting the start of exon2 of *SLAMF6* in Jurkat cells. The frameshift mutation generated is highlighted (red). Only mRNA in which the sequence encoding aa 17-65 is skipped yields a viable protein. **b**, An illustration of the antibodies used for both flow cytometry and western experiments. Anti-SLAMF6 (NT7) targeting the SLAMF6 V-domain, only detecting the canonical isoform, and anti-SLAMF6 (AF1908) targeting SLAMF6 C-domain. **c**, SLAMF6 canonical isoform expression in WT Jurkat cells and Jurkat-SLAMF6^Δ17-65^ cells stained with anti-SLAMF6 (NT7) targeting the SLAMF6 V-domain. **d**, Immunoblot of SLAMF6 in WT Jurkat cells and Jurkat-SLAMF6^Δ17-65^ cells using anti-SLAMF6 (AF1908) targeting SLAMF6 C-domain. **e**, SLAMF6^Δ17-65^/ SLAMF6^ΔExon2^ isoform expression in WT Jurkat cells and Jurkat-SLAMF6^Δ17-65^ cells stained with anti-SLAMF6 (AF1908) targeting the SLAMF6 C-domain. **f**, Quantitative RT-PCR for *IL-2* in WT Jurkat versus Jurkat-SLAMF6^Δ17-65^. Data were normalized to *HPRT* expression for each cell type and for time point 0. **g**, ELISA for IL-2 produced in WT Jurkat, Jurkat-SLAMF6^Δ17-65^, and Jurkat-SLAMF6 KO cells activated for 48 h with PMA-ionomycin. **h**, ELISA for IL-2 in WT Jurkat and Jurkat-SLAMF6^Δ17-65^ cells electroporated with siRNA targeting SLAMF6 and activated for 48 h. One-way ANOVA. *, P < 0.05, **, P < 0.01, ***, P < 0.001.

Using RNA sequencing (detailed in the next section), we identified the skipping of junction 2, which is the splicing site for the canonical SLAMF6. In accordance, we observed an increased prevalence of the alternative junction, joining together the codons encoding for amino-acids 16 and 66 (Sashimi plot, Supplementary figure 3a). Quantitative analysis of SLAMF6 isoforms expression in the cells showed that following CRISPR-Cas 9 editing, the transcript coding for the canonical SLAMF6 is decreased, probably due to the process of nonsense-mediated decay^24^ (Supplementary figure 3b).

Using antibodies targeting the variable and constant regions of SLAMF6 (Figure 4b), we showed that the edited Jurkat cells did not express the canonical SLAMF6 receptor but had detectable levels of SLAMF6^Δ17-65^ protein (Figure 4c,d), which was expressed on the cell membrane (Figure 4e). The expression of other SLAM family receptors and ligands remained unchanged (Supplementary figure 3c).

After verifying that SLAMF6^Δ17-65^ exists as a viable protein on the cell surface, we evaluated the functional capacity of Jurkat-SLAMF6^Δ17-65^ cells. Cytokine secretion, measured using IL-2 transcript and ELISA following 2-day activation, showed a three-fold increase in Jurkat-SLAMF6^Δ17-65^ cells compared with the parental line (Figure 4f,g and Supplementary figure 3d,e). Of note, Jurkat cells missing all SLAMF6 isoforms did not show an increased level of IL-2, indicating that the splice isoform was acting on its own and not via the absence of the canonical SLAMF6 inhibitory receptor. The positive effect of SLAMF6^Δ17-65^ was obliterated by siRNA designed to knock down he gene (Figure 4h and Supplementary figure 3f). While activated Jurkat cells do not secrete detectable levels of IFN-γ, Jurkat-SLAMF6^Δ17-65^ cells did produce IFN-γ (Supplementary figure 3g), attesting for polyfunctionality.

### Activated SLAMF6^Δ17-65^ Jurkat cells acquire a cytotoxic gene expression program

To generate a comprehensive transcription map of Jurkat-SLAMF6^Δ17-65^ cells, longitudinal RNA sequencing was performed following activation with PMA and ionomycin. In a principal-component analysis to map cell populations in two-dimensional space (Figure 5b), WT and Jurkat-SLAMF6^Δ17-65^ cells were observed to cluster closely before activation, but were substantially separated at all time points after activation. We noted that the splice isoform induced some distinct traits in the lymphocytes (Figure 5a). Most prominent among them was the expression of *SLAMF7*, which shortly after activation was highly and constitutively expressed in the Jurkat-SLAMF6^Δ17-65^ cells, at all time points sampled; at several time points *SLAMF1* was also expressed. SLAMF7 is typically expressed in activated NK and CD8 T cells, but not in Jurkat or CD4 T cells. The *de novo* transcription of SLAMF7 was accompanied by a major rise in *GZMB and NKG2D (KLRK1)*, delineating a transition from a CD4 to an NK/CD8 profile. A high NKG2D expression at the protein level was also noted in 72h-activated Jurkat-SLAMF6^Δ17-65^ cells compared to WT (Figure 5c). When combined with the additional increased expression of *TNF, IFNG, IL2, CD30LG*, and *LIGHT*, this profile gives rise to an improved cytotoxic score of the Jurkat-SLAMF6^Δ17-65^ cells^25^ (Figure 5d). Higher levels of chemokines *CCL3* and *CCL5* suggest a favorable migration capacity (raw data, not shown).

**Figure 5:**
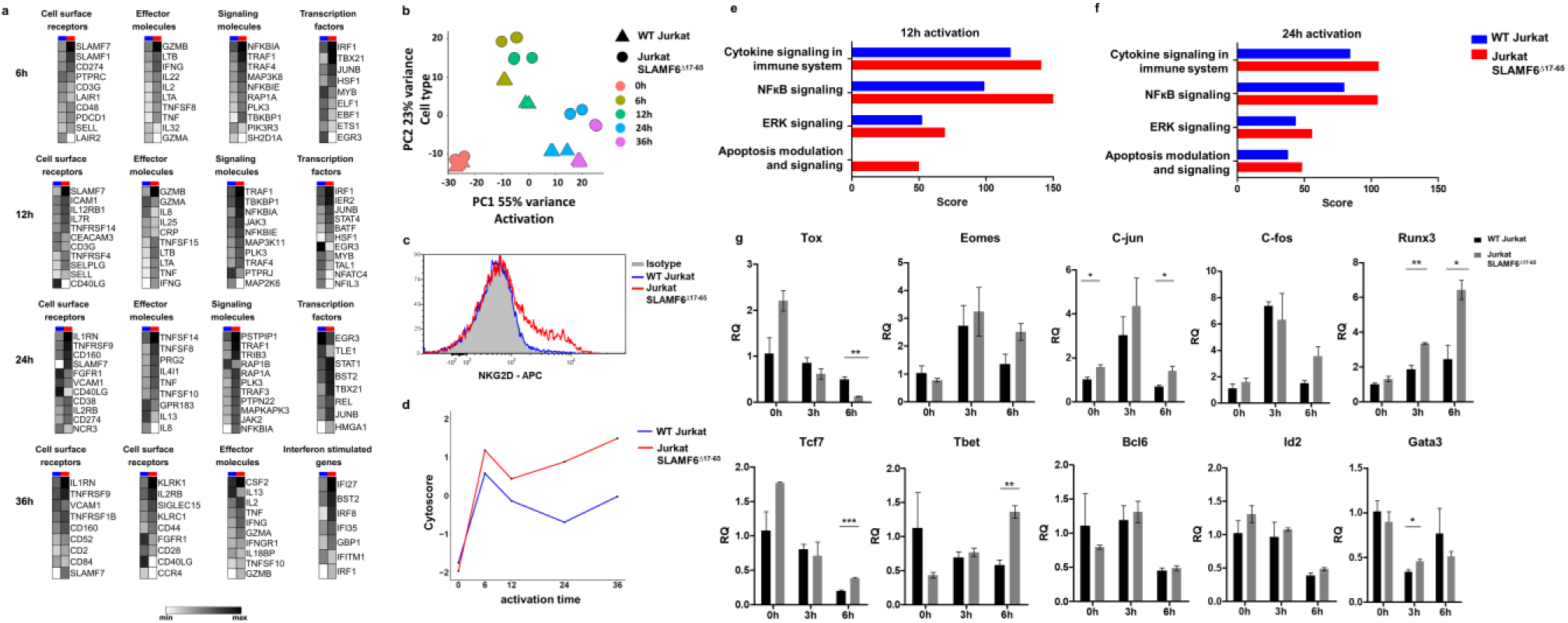
Activated SLAMF6^Δ17-65^ Jurkat cells acquire a cytotoxic gene expression program. **a**, heatmap illustrating the average transcript expression of the indicated genes during activation. Color scale represents row means. **b**, Principal component analysis (PCA) of the RNA sequencing data that characterizes the trends exhibited by the expression profiles of WT Jurkat (triangle) and Jurkat-SLAMF6^Δ17-65^ (circle) cells during activation. Each dot represents a sample, and each color represents the time of activation. **c**, NKG2D expression in WT Jurkat and Jurkat-SLAMF6^Δ17-65^ cells following 72h of activation. **d**, Cytotoxic score during activation, comparing WT Jurkat and Jurkat-SLAMF6^Δ17-65^ cells. **e-f**, GO enrichment of 4 selected pathways comparing WT Jurkat and Jurkat-SLAMF6^Δ17-65^ cells following 12 h **(e)** and 24 h **(f)** of activation. **g**, Quantitative RT-PCR for transcription factor expression in WT Jurkat and Jurkat-SLAMF6^Δ17-65^ cells performed 6 h following activation. Data were normalized to *HPRT* expression for each cell type and WT Jurkat at 0 h. One-way ANOVA. *, P < 0.05, **, P < 0.01, ***, P < 0.001.

In parallel with the acquisition of a cytotoxic CD8-like profile, the Jurkat-SLAMF6^Δ17-65^ cells completely lost expression of *CD40LG*, which was constitutively expressed in the WT cells. CD40LG takes part in the CD4 T cell-regulated B cell response. Its loss adds to the depth of the change resulting from SLAMF6 alternative splicing, which affects a variety of functions that are not strictly co-stimulatory.

Analysis of differentially expressed genes (DEGs) coupled with GO term enrichment analysis was used to compare transcriptional programs of the lymphocytes (Figure 5e,f). SLAMF6^Δ17-65^ cells showed more substantial usage of NFkB and ERK signaling with a higher susceptibility to apoptosis than WT Jurkat cells. Lastly, Jurkat-SLAMF6^Δ17-65^ cells had an intriguing transcription factor profile, which was distinct from that of the parental line (Figure 5a,g). Levels of *Tbet, Runx3, C-jun*, and *Tcf7*, which typify effector T cell subsets, were significantly higher in activated Jurkat-SLAMF6^Δ17-65^ cells than in WT cells. In comparison, already at six hours the parental line expressed high levels of *EGR3*, an immune response-restraining transcription factor, and this factor remained highly expressed until 24h, *EGR3* had hardly any expression in Jurkat-SLAMF6^Δ17-65^ cells in the early time points. *Tox*, a key regulator of the exhuasted state, decreased in expression in both cell populations, but the reduction was more substantial for Jurkat-SLAMF6^Δ17-65^ cells. In summary, the deletion in CD4 Jurkat cells of full-length SLAMF6, together with preferential expression of the SLAMF6 isoforms SLAMF6^Δ17-65^/SLAMF6Δ^exon2^, induces a cytotoxic gene transcription program characterized by *Tbet* expression and the elimination of *EGR3*. Thus, an array of cytokines and cytolytic genes are overexpressed in the modified T cells.

### SLAM-associated protein (SAP) is reduced in activated SLAMF6^Δ17-65^ Jurkat cells while SHP1 is required for their increased cytokine production

The ITSMs of SLAMF6 serve as docking sites on the cytoplasmic tail for the SH2 domain-containing adaptor SAP, which couples to all SFRs^26^. SAP plays a central role in the collaboration of B cell and T cells and is considered critical for their activation^27^. It was therefore surprising to find that the SAP transcript was reduced in Jurkat-SLAMF6^Δ17-65^ cells (Figure 6a, at 6 h of activation, p=0.007, and at 36 h of activation, p<0.0001), with parallel results at the protein level (Figure 6b). To investigate the role of SAP in the Jurkat-SLAMF6^Δ17-65^ cells, we knocked down the residual SAP with siRNA and observed that IL-2 secretion increased further (Figure 6c,d). XLP (X-linked lymphoproliferative disease) is a fatal immune dysregulation syndrome, caused by a mutation creating non functioning SAP and characterized by unremitting Epstein-Barr virus (EBV) infections, dysgammaglobulinemia, and lymphoma^28^. PBMCs obtained from two XLP patients with SAP mutations displayed a very high IFN-γ secretion of 9000 pg/ml, compared to 250 pg/ml secreted by PBMCs from healthy donors tested in the same experiment and under the same activation conditions (Figure 6e). Indirectly, the XLP patient data hints that SAP depletion augments cytokine secretion, as was seen in the SLAMF6^Δ17-65^ cells.

**Figure 6:**
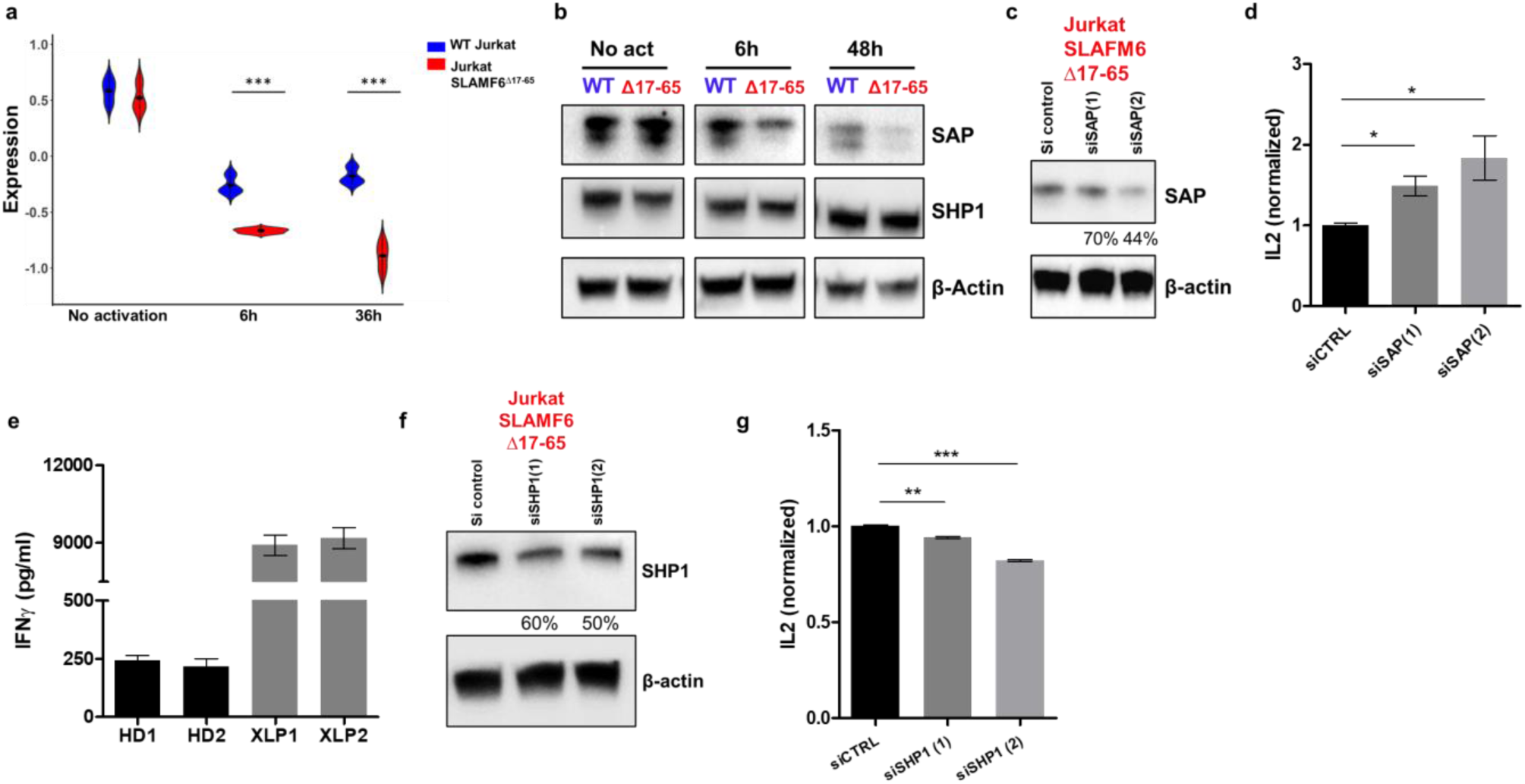
SLAM-associated protein (SAP) is reduced in activated Jurkat-SLAMF6^Δ17-65^ cells, and SHP1 is required for their increased cytokine production. **a**, Expression of *SH2D1A* among WT Jurkat and Jurkat-SLAMF6^Δ17-65^ cells at 3 time points during activation, presented as DESeq2 normalized in violin plots. **b**, Immunoblot for SHP1 and SAP during activation of WT Jurkat and Jurkat-SLAMF6^Δ17-65^ cells activated for 48 h. **c-d**, IL-2 secretion by SAP knocked-down Jurkat-SLAMF6^Δ17-65^ cells. **c**, Immunoblot analysis following siRNAs targeting SAP. **d**, IL-2 secretion 48 h after activation. **e**, IFN-γ secretion by PBMCs from two XLP patients activated overnight with plate-bound anti-CD3 antibody. HD, healthy donor. **f-g**, IL-2 secretion by SHP1 knocked-down Jurkat-SLAMF6^Δ17-65^ cells. **e**, Immunoblot analysis following siRNAs targeting SHP1. **f**, IL-2 secretion 48 h after activation. CTRL, control siRNA.

In contrast, silencing SHP-1 curtailed IL-2 secretion (Figure 6f,g), suggesting that SHP-1 is necessary for the agonistic activity of the SLAMF6^Δ17-65^ isoform. Coupling of SHP-1, a protein phosphatase, to SLAMF6 has previously been associated with reducing T cell help to B cells^29^. Its effect on the SLAMF6^Δ17-65^ cells draws attention to the dependence of signal-propagating molecules on both cell-type and context. Thus, although the cytoplasmic tail of SLAMF6^Δ17-65^ remains unchanged, its modulatory role reduces SAP expression and requires SHP-1 collaboration, generating outcomes opposite to those described for full-length SLAMF6.

### SLAMF6 splicing pattern changes during treatment of metastatic melanoma patients with checkpoint inhibitors

Immune checkpoint blockade (i.e. monoclonal antibodies targeting CTLA-4 and PD-1) that reverses T cell dysfunction changed the prospects for melanoma patients. PD-1 blockers, in particular, have had a significant impact on T cell functionality^30,31^. We, therefore, evaluated how ICB therapy would affect the ratio of the SLAMF6 isoforms. To address this question, PBMCs were obtained from seven metastatic melanoma patients before and during ICB therapy. CD8 subsets were separated, and *SLAMF6* isoforms were detected using RT-PCR (Figure 7a). In healthy donors, isoform expression was measured at several time points and found to be consistently unchanged (data not shown). In four melanoma patients, no change in isoform expression was noted after ICB treatment, with one patient achieving a complete response. In the other three patients, a higher expression of *SLAMF6*^*Δ17-65*^ was recorded after treatment, in the effector and effector memory (EM) CD8 subsets (Figure 7b-c). Interestingly, these three patients with higher *SLAMF6*^*Δ17-65*^ expression suffered severe adverse effects that necessitated treatment disruption: one had grade IV liver toxicity on a combination of CTLA4 and PD-1 inhibitors (the patient is now in complete remission), and two had grade III diarrhea (one attained partial response but then progressed; the other received preventive treatment). Of these, patient 3 (Figure 7d), who received a combination of CTLA4 and PD-1 inhibitors, was switched to PD-1 monotherapy due to the toxicity. The change in treatment was associated with a reciprocal decrease in *SLAMF6*^*Δ17-65*^ in his effector CD8 T cell populations. The connection between higher *SLAMF6*^*Δ17-65*^ expression and treatment toxicity may suggest that ICB induces splicing events in lymphocytes that are part of their effect on the immune system.

**Figure 7:**
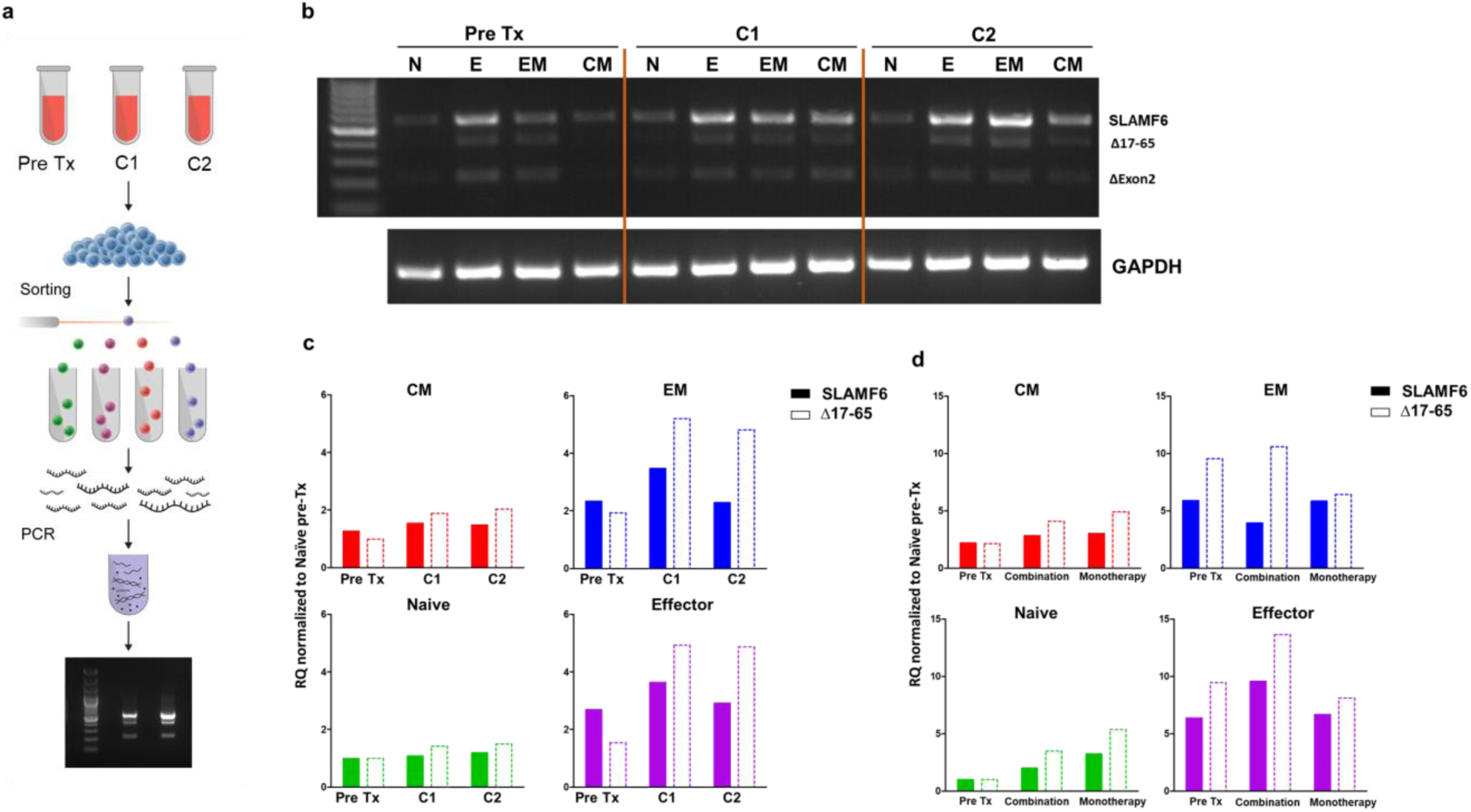
SLAMF6 splicing pattern changes during treatment of metastatic melanoma patients with checkpoint inhibitors. **a**, Blood samples from metastatic melanoma patients were taken at three time points before starting immune checkpoint blockade (ICB) treatment and twice during the treatment, three months apart. Samples from a healthy donor were also taken three months apart. CD8 T cells from each sample were separated into subsets as described in supplementary figure 1B. For each subset, RNA was extracted, and quantitative RT-PCR for *SLAMF6* isoforms was performed. Data were normalized to *HPRT* expression for each subset. For each isoform, data were normalized to the value of naϊve cells at the 1^st^ time point. **b**, Representative agarose gel showing RNA expression of *SLAMF6* isoforms for one of the patients during the ICB treatment. **c**, Quantitative RT-PCR for *SLAMF6* isoforms at each time point for patient 1. Each T cell subset is shown separately. **d**, Quantitative RT-PCR for *SLAMF6* isoforms at each time point for patient 3. This patient had a change of treatment regimen to a less toxic one. N, naϊve. E, effector. EM, effector memory, CM, central memory.

### SLAMF6 splice switching with antisense oligonucleotides leads to improved anti-tumor T cell response

To investigate the role of SLAMF6^Δ17-65^ in healthy T cells, we developed a method to modify SLAMF6 splicing in genetically non-manipulated cells. ASOs, which are short, single-stranded DNA molecules that interact with mRNA, were used as the molecular modifier for this purpose. ASOs were initially developed to prevent the translation of genes that lead to pathological RNA or protein gain-of-function, such as occurs in muscular degenerative diseases^32^. An ASO that tightly anneals to an exonic target may prevent the splicing machinery from joining together two exons while removing the intervening intron^33^ (Figure 8a). The SLAMF6^Δ17-65^ isoform is typically expressed in T cells at a very low level, compared with the canonical isoform (Figure 1d). To augment the proportion of the SLAMF6^Δ17-65^ isoform, we designed a splice-switching ASO with RNA oligonucleotides synthesized with a full (all nucleotides) phosphorothioate (PS) backbone and in which each ribose 2′-hydroxyl was modified to 2′-methoxyethyl (2′-MOE). After electroporation into Jurkat cells, the designed ASO changed the splicing of *SLAMF6* in favor of *SLAMF6*^*Δ17-65*^ (Figure 8b, Supplementary figure 4a,b). Higher IL-2 secretion was induced by this change (Figure 8c). Transcription factor expression in the ASO-transfected cells was almost identical to that found in the Jurkat-SLAMF6^Δ17-65^ cells, characterized by increased *Tbet, C-jun, and Runx3*, and decreased *Tox* levels (Figure 8d). In primary T cells, activated ASO-modified lymphocytes showed higher IFN-γ secretion compared to non-modified cells (Figure 8e). Finally, we implanted human melanoma cells mixed with splicing-modified and non-modified cognate TILs (209*TIL*) into nude athymic mice (Figure 8f,g). Mice injected with 209*TIL* that underwent ASO-induced SLAMF6^Δ17-65^ preferential expression showed improved tumor control (Figure 8h), maintaining their superior effect for as long as 30 days after tumor injection (Figure 8i,j). These results demonstrate the feasibility of using antisense oligonucleotides in adoptive cell therapy and emphasize the power of splicing modification to produce T cells with improved anti-tumor characteristics.

**Figure 8:**
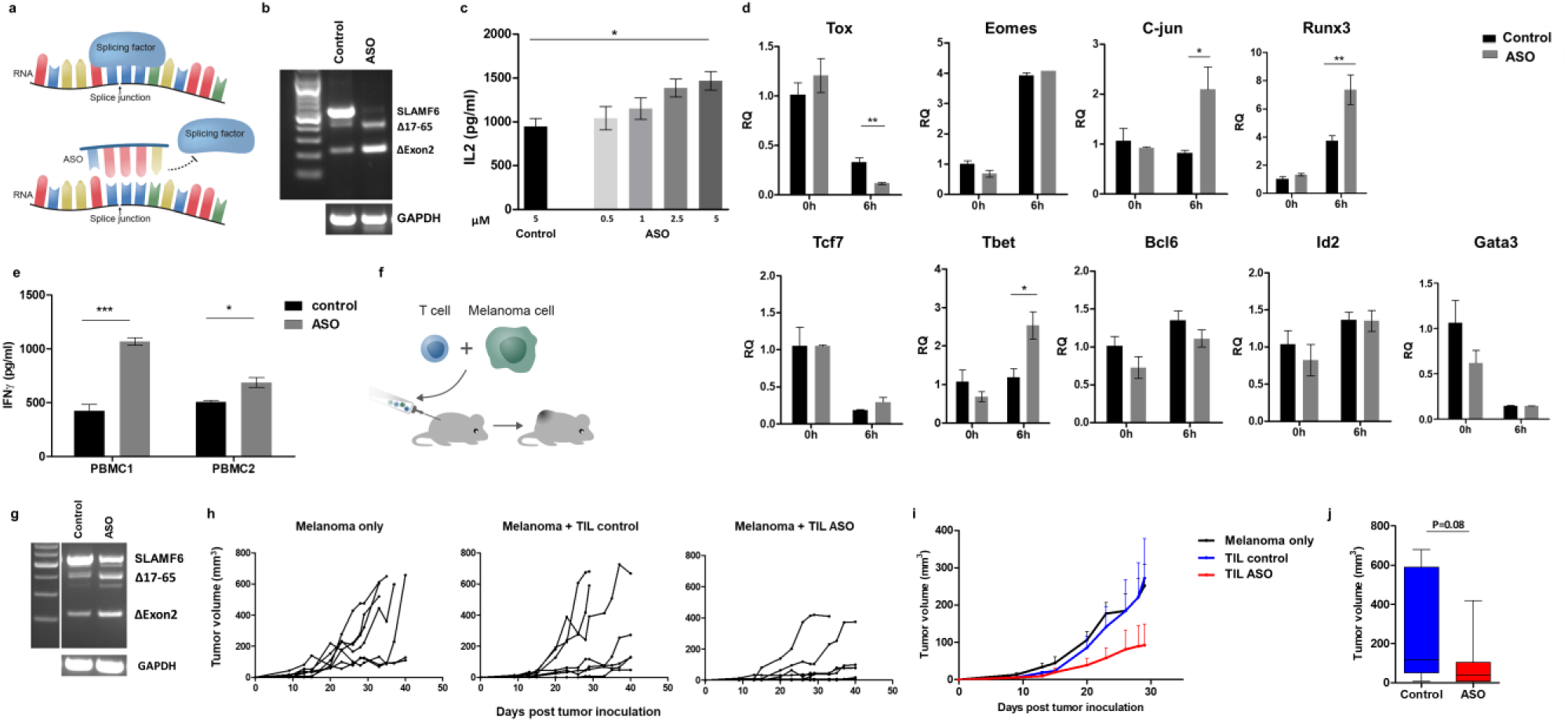
SLAMF6 splice switching with antisense oligonucleotides leads to improved anti-tumor T cell response. **a**, Schematic diagram of the antisense-oligos (ASOs) targeting the splicing site of SLAMF6 in exon2. **b**, RNA expression of *SLAMF6* isoforms in WT Jurkat cells electroporated with scrambled ASO or ASO targeting the splice site. **c**, ELISA for IL-2 in WT Jurkat cells activated for 48 h after electroporation with scrambled ASO or increasing concentration of ASO targeting the splice site. **d**, Quantitative RT-PCR of WT Jurkat cells electroporated with ASO or control ASO and activated for 6 h. Data were normalized to *HPRT* expression for each manipulation. Further normalization for each transcript comparing to control ASO (0 h). **e**, ELISA for IFN-γ in anti-CD3 plate-bound activated PBMCs, 24h after electroporation with ASO, or control ASO. **f-j**, TILs were electroporated with ASO or control ASO. Twenty-four h after electroporation, the cells were washed and mixed at a 1:1 ratio with 526*mel* cells and immediately injected subcutaneously into the back of nude (athymic Foxn1-/-) mice (N=7). **f**, Schematic diagram showing the experimental layout. **g**, RNA expression of *SLAMF6* isoforms in TILs electroporated with scrambled ASOs or ASOs targeting the splice site. **h**, Spider plot showing tumor volume [calculated as L (length) x W (width)^2^ x 0.5]. **i**, Tumor volume (Mean+SEM) until day 29, when the first mouse was sacrificed. **j**, Mean+SEM tumor volume on day 29. One-way ANOVA. *, P < 0.05, ***, P < 0.001.

## Discussion

Alternative splicing (AS) is a process by which one gene encodes more than one protein by removing and rejoining segments of exons or retaining parts of introns. The outcome of AS is a set of proteins derived from the same DNA sequence but having diverse and even contradictory functions. In cancer research, thousands of splice variants have been identified in the last decade that result from mutations in splice sites and regulatory elements, or from an erratic splicing process ^34,35^. We investigated the AS of the SLAMF6 immune receptor, to evaluate the role of this process in the immune response to cancer.

AS is a mechanism that yields protein diversity and also a dynamic process that shapes variant protein repertoires across cell types, origins, and states. Using the Immunological Genome Project transcriptional data, Ergun et al. showed that over 70% of the genes expressed in immune cells undergo AS, and that exon patterns cluster according to lineage differentiation and cell cycling^36^. The type of stimuli dictates varied isoform usage in activated monocytes^37^ and splicing pattern shifts in activated macrophages^38^, dendritic cells exposed to influenza^39^, and T and B lymphocytes^36^. Some genes co-expressed in B and T cells had different isoform ratios in these populations. The Porgador group has reported that decidual NK cells in healthy pregnancies display diverse NK receptor splicing compared to cells isolated from miscarriages^40^. This report is among the earliest to link AS of immune receptors to a change of function.

The well-known example of AS that determines a functional state, is surface phosphatase CD45. The classical marker of naϊve T cells, CD45RA, is the longer variant of CD45, which de-phosphorylates inhibitory tyrosine kinases at a slower rate than CD45RO, an isoform lacking exons 4, 5, and 6. In fact, CD45RO defines effector and memory T lymphocytes. The fast activation of the latter subsets is attributed to loss of the CD45RO exon and consequent enhanced function^41^.

Here we report that the alternative splicing of an immune receptor gene, *SLAMF6*, generates two opposing modulatory activities. The mode of action of the shorter isoform differs from the canonical variant at all levels investigated, including transcriptional events, signal generation, and modulatory function. The canonical SLAMF6 is a constitutively expressed self-binding receptor whose expression increases in activated T cells and which inhibits their function, acting as a type of rheostat. The shorter variant, SLAMF6^Δ17-65^, acts as a co-stimulatory receptor that strongly enhances T cell activation and, in our model system, augments an anti-melanoma response. The change in function that the splicing process induces in SLAMF6 is profound and perhaps unique. The transcriptional landscape of SLAMF6^Δ17-65^ is governed by the *SLAMF7, GZMB*, and *NKG2D* genes, which confer cytotoxic activity, and by the transcription factors *Tbet* and *Runx3*. These features are encountered in helper and exhausted progenitor phenotypes, cell states that develop in activated T cells and that characterize subsets critical for an effective immune response^42,43,44,45,46^. Thus, while SLAMF6 is an identifier of exhausted T cell precursors, it is also a key player in heightened T cell functionality. The work presented here suggests that SLAMF6^Δ17-65^, and not the canonical SLAMF6, is the driver of the effector traits.

The patient data (figure 7), which shows that the SLAMF6 splicing ratio may shift during treatment, supports our conclusion that SLAMF6^Δ17-65^ is involved in the effect of PD-1 and CTLA4 blocking antibodies. The increase in the expression of *SLAMF6*^*Δ17-65*^ transcript was greater with CTLA4 blockade and during severe toxicity than with PD-1 blockade alone. The expression of this agonistic isoform decreased during corticosteroid treatment. Finally, the therapeutic power of SLAMF6^Δ17-65^ was demonstrated using splice-switched melanoma cognate TILs (figure 8), which were superior in their capacity to inhibit tumor growth in nude mice. Interestingly, full-length SLAMF6 was not abolished in the adoptively transferred T cells. Hence, it is most plausible that the increase in SLAMF6^Δ17-65^ protein was responsible for the improved activity rather than a knockdown effect of SLAMF6.

The signal generation following SLAMF6 transactivation depends on the recruitment of SAP, which is named for its association with the SLAM family (SLAM Associated Protein). However, SAP also adapts to PD-1, which in addition to its immune tyrosine inhibitory motif, contains a switch signaling motif^47,48^. The depletion of SAP protein noted in Jurkat-SLAMF6^Δ17-65^ cells is central to its agonistic effect but unrelated to its cytoplasmic sequence, which was not altered. The absence of functional SAP led to a considerable amount of IFN-γ secretion in T cells from XLP patients, and likewise, SAP silencing enhanced IL-2 secretion in Jurkat-SLAMF6^Δ17-65^ cells. Therefore, it is reasonable to assume that SAP recruitment is inseparable from the SLAMF6 rheostatic effect. SAP depletion may hamper other regulatory SFRs, mainly SLAMF4^49^. In contrast, SHP-1, which presumably displaces SAP, participates in the agonistic effect, as shown in the silencing experiments (figure 6g). Since inhibitory tyrosine kinases are among the targets of SHP-1, we propose that by inactivating them, SHP-1 mediates the SLAMF6^Δ17-65^ agonistic effect. This mechanism was attributed to NK cell education via SLAMF6 against hematopoietic targets^50^. Altogether, SLAMF6 splice-switched T cells generate a strong, integrated initial stimulus. The cohesive molecular program that ensues initiates an array of cytotoxicity genes and directs the T cells towards a cytotoxic phenotype, even if they originate in the CD4 lineage, as the Jurkat cell data shows.

The functional advantage that heightened SLAMF6^Δ17-65^ confers on anti-tumor CD8 T cells is an asset for adoptive cell therapies. Splice-switching oligonucleotide drugs are attracting attention based on success in clinical trials for Duchenne muscular dystrophy (DMD) and spinal muscular atrophy (SMA)^51,52^. In DMD, interfering with the normal splicing process induces skipping of an exon containing a frameshift deletion, restores the mRNA reading frame, and produces an internally deleted protein that improves muscle function^53^. Chemical modification of nucleic acids, with improved base-pairing affinity and specificity as well as increased resistance to nucleases, has increased the utility of ASO in the clinic^54^. While a significant problem in genetic diseases is penetrance to the affected tissues, *ex vivo* modification of T cells prior to their adoptive transfer avoids most of these drawbacks. The encouraging tumor growth inhibition achieved by the anti-melanoma TILs in which the ratio between SLAMF6 isoforms had been switched highlights the potential of this approach as a novel method for adoptive cell therapy (figure 8h).

In summary, the immune system faces a significant challenge in confronting disease, as it must mount a rapid, effective response while maintaining homeostatic regulation to prevent collateral damage. The example of SLAMF6 demonstrates how one receptor gene achieves this balance. SLAMF6 encodes two tightly coordinated modulatory isoforms that act in opposite ways and change their ratio in a context-dependent manner. The danger entwined in the agonistic SLAMF6^Δ17-65^ isoform is kept in check by the co-expression of the canonical inhibitory receptor. This balance, however, may be disrupted by therapeutic intervention, as seen with the rise of the shorter transcript in patients experiencing ICB toxicity. ASOs that switch splicing to favor the SLAMF6^Δ17-65^ isoform yield T cells with augmented tumor control. Judicious use of such targeted ASOs may therefore constitute a new strategy to improve cell-based immunotherapies.

## Materials and methods

### Plasmids

#### pCDNA3.1 –DYK-slamf6 isoforms

pcDNA3.1+/C-(K)DYK-slamf6 transcript isoforms were purchased from Genscript (OHu04772, OHu04774, OHu4776, and the empty vector).

#### pLL3.7-FLAG-slamf6 isoforms

Plasmids for each SLAMF6 isoform expression were produced according to the following procedure: the Flag tag was inserted by fusion PCR between the signal sequence and the rest of the ORF of the SLAMF6 isoform. The two fused PCR sequences were amplified with the following pairs of primers:

VAR-5: TATATTCTAGACCATGTTGTGGCTGTTCCAATCG

VAR1-3: TATATGAATTCTTACACGACATTGTCAAGGGC

Each PCR sequence was cut with XbaI + EcoRI and then cloned into NheI / EcoRI-cut lenti vector pLL3.7. Sequencing results indicated that the genes were cloned as designed, and no non-specific mutations were present.

For the control empty vector, the pLL3.7 vector was cut with NheI + EcoRI and treated with Klenow fragment enzyme followed by self-ligation. The sequencing result indicated that the EGFP from the vector was deleted.

#### pSpCas9(BB)-2A-GFP CRISPR plasmids

pSpCas9(BB)-2A-GFP plasmids for SLAMF6 KO were produced according to the Zhang protocol^55^, using the following guides:

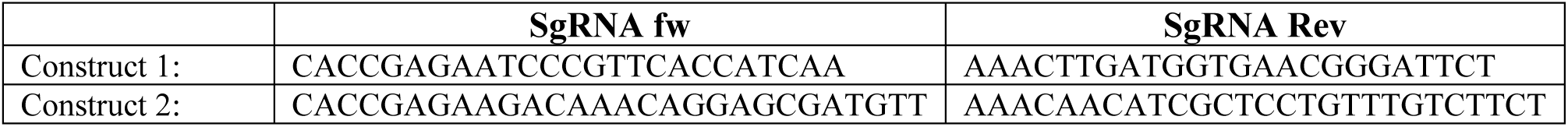

### Cells

#### Melanoma cells

The cell line 526*mel* (HLA-A2+/MART-1+gp100+) was a gift from M. Parkhurst (Surgery Branch, NCI, NIH). The cells were cultured in RPMI 1640 supplemented with 10% heat-inactivated fetal calf serum (FCS), 2 mmol/L L-glutamine, and combined antibiotics (all from ThermoFisher Scientific, Massachusetts, USA). All lines are regularly tested and were mycoplasma free.

#### Aberrant slamf6 splice variant expression on melanoma cells

526*mel* human melanoma cell lines were transfected with the pcDNA3.1**+**/C-(K)DYK-neomycin-slamf6 transcript isoforms using lipofectamine 2000 (ThermoFisher). Neomycin resistant melanoma cells were subcloned, and the stably transfected cells were labeled with anti-SLAMF6 Ab and sorted in an ARIA-III sorter. Cells transfected with the empty vector were sub-cloned using neomycin resistance.

#### Hel293 cells

The Hek293 cell line was purchased from ATCC. The cells were cultured in DMEM supplemented with 10% heat-inactivated fetal calf serum (FCS), 2 mmol/L L-glutamine, and combined antibiotics (all from ThermoFisher Scientific). The line is regularly tested and was mycoplasma free.

#### Aberrant slamf6 splice variant expression on Hek293 cells: Hek293

cells were transfected with the pLL3.7-FLAG-slamf6 transcript isoforms or the empty control vector using lipofectamine 2000 (ThermoFisher). Neomycin resistant melanoma cells were subcloned, and the stably transfected cells were labeled with anti-SLAMF6 Ab and sorted in an ARIA-III sorter. Cells transfected with the empty vector were sub-cloned using neomycin resistance.

#### Jurkat cells

The Jurkat cell line was purchased from ATCC. The cells were cultured in RPMI 1640 supplemented with 10% heat-inactivated fetal calf serum (FCS), 2 mmol/L L-glutamine, and combined antibiotics (all from ThermoFisher Scientific). The line is regularly tested and was mycoplasma free.

#### SLAMF6 KO Jurkat cells

SLAMF6 KO Jurkat cells were generated using CRISPR-Cas mediated genome editing, using the px330 plasmid with the sgRNA described above. Jurkat cells were washed twice in RPMI medium without supplemets and resuspended in 10×10^6^ cells/ml RPMI. 5×10^6^ Jurkat cells with 5μg SLAMF6-CRISPR plasmid were electroporated in Biorad 0.4cm cuvettes using ECM 630 Electro Cell manipulator (BTX Harvard apparatus) at 260V, 975μF, 1575Ω. After electroporation, cells were immediately seeded into complete RPMI medium. 48h after transfection, cells expressing GFP were selected by sorting (ARIA-III Sorter). Cells lacking human SLAMF6 were selected by single-cell sorting in 96 wells using anti-SLAMF6 Ab, and cultured until single-cell colonies grew. The genomic DNA from each colony was sequenced to confirm the mutation.

#### T2 cells

The T2 cell line was a kind gift from Lea Eisenbach, Weizmann Institute, Israel. The cells were cultured in RPMI 1640 supplemented with 10% heat-inactivated fetal calf serum (FCS), 2 mmol/L L-glutamine, and combined antibiotics (all from ThermoFisher Scientific). The line is regularly tested and was mycoplasma free.

#### SLAMF6 isoforms perturbated T2 cells

T2 with perturbated SLAMF6 isoforms cells were generated using CRISPR-Cas mediated genome editing, using the px330 plasmid with the sgRNA, as described above. T2 cells were washed twice in RPMI medium without supplements and resuspended in 10×10^6^ cells/ml RPMI. 5×10^6^ T2 cells with 5μg SLAMF6-CRISPR plasmid were electroporated in Biorad 0.4cm cuvettes using ECM 630 Electro Cell manipulator (BTX Harvard apparatus) at 260V, 975μF, 1575Ω. After electroporation, cells were immediately seeded into complete RPMI medium. Forty eight hours after transfection, cells expressing GFP were selected by sorting (ARIA-III Sorter). Single cells were sorted in 96 wells and cultured until single-cell colonies grew. The genomic DNA from each colony was sequenced to validate the mutation.

#### Peripheral blood mononuclear cells (PBMC)

were purified from buffy coats of healthy donors (Hadassah Blood Bank) or purified from samples taken from patients undergoing ICB treatment (Helsinki approval 383-23.12.05). Blood samples from metastatic melanoma patients were obtained at three time points: once before starting immune checkpoint blockade (ICB) treatment and twice during the treatment, three months apart. The cells were sorted into four CD8+ subpopulations using an ARIA-III sorter, based on staining for CD8, CCR7, and CD45-RA markers. For each subset, RNA was extracted, and quantitative RT-PCR for SLAMF6 isoforms was performed. All patients gave their informed consent to participate in this study.

#### Tumor-infiltrating lymphocytes (TILs)

Fresh tumor specimens taken from resected metastases of melanoma patients were used to release TILs using a microculture assay (Lotem *et al*., 2002). Human lymphocytes were cultured in complete medium (CM) consisting of RPMI 1640 supplemented with 10% heat-inactivated human AB serum, 2 mmol/l L-glutamine, 1 mmol/l sodium pyruvate, 1% nonessential amino acids, 25 mmol/l HEPES (pH 7.4), 50 μmol/l 2-ME, and combined antibiotics (all from Invitrogen Life Technologies). CM was supplemented with 6000 IU/ml recombinant human IL-2 (rhIL-2, Chiron).

#### Cloning of peptide-specific TILs

On day 14 after tumor-infiltrating lymphocyte (TIL) culture initiation, lymphocytes were washed with PBS, resuspended in PBS supplemented with 0.5% BSA, and stained with FITC-conjugated HLA-A*0201*/*MART-126–35 or HLA-A*0201*/*gp100209-217 dextramer (Immudex) for 30 minutes at 4°C. Lymphocytes were then incubated with allophycocyanin-conjugated mouse anti-human CD8 (eBioscience) for an additional 30 minutes at 4°C and washed. CD8+ lymphocytes, positively stained by dextramer, were sorted with a BD FACS-Aria and directly cloned at one or two cells per well in 96-well plates in the presence of anti-CD3 (30 ng*/*mL, eBioscience), rhIL-2 (6,000 IU/mL), and 4 Gy-irradiated 5×10^4^ allogeneic PBMCs as feeder cells. Five days later, rhIL2 (6,000 IU/mL) was added and renewed every two days. On day 14, the clones were assayed for IFN-γ secretion in a peptide-specific manner following their coincubation with T2 cells pulsed with MART-1_26–35_ or gp100_209-217_ [both commercially synthesized and purified (>95%) by reverse-phase HPLC by Biomer Technology] by ELISA (R&D Systems). The MART-1_26– 35_ or gp100_209-217_–reactive clones were further expanded in a second-round exposure to anti-CD3 (30 ng/mL), and rhIL2 (6,000 IU/mL) in the presence of a 50-fold excess of irradiated feeder cells.

### Antibodies

For flow cytometry, cells were labeled with the following reagents: anti-SLAMF6 (NT-7), anti-CD45RA (HI100), anti-CCR7 (G043H7), anti-CD62L (DREG-56), anti-CD8 (RPA-T8), anti-TNFα (Mab11), anti-CD244 (C1.7), anti-CD48 (BJ40), anti-CD229 (HLy-9.1.25), anti-CD84 (CD84.1.21) all obtained from Biolegend (San Diego, CA, USA). Anti-SLAMF6 (REA) was obtained from Miltenyi Biotec (Bergisch Gladbach, Germany). Anti-IFN-γ (4S.B3) was obtained from Biogems (Westlake Village, CA, USA), anti-FLAG (F3165) obtained from Sigma-Aldrich and anti-CD150 (A12(7D4)) from eBioscience (California, USA). For immunoblotting, proteins were detected with anti-SLAMF6 (Goat, AF1908, R&D Systems, Minneapolis, MN, USA), anti-SAP (Rat, 1A9, Biolegend), anti-SHP1 (Rabbit, Y476, Abcam, Cambridge, UK), and anti-β actin (Mouse, sc-47778, Santa Cruz Biotechnology, Texas, USA).

seSLAMF6, a soluble polypeptide consisting of the ectodomain sequence of SLAMF6, was purchased from Novoprotein^23^. seSLAMF6^Δ17-65^, a soluble polypeptide consisting of the ectodomain sequence of SLAMF6^Δ17- 65^ isoform, was purchased from Bon Opus Biosciences.

### Mice

Nude (athymic Foxn1^-/-^) mice were purchased from Harlan Laboratories.

### In vitro assays

#### RT-PCR and qRT-PCR

RNA was isolated from cells using the GenElute Mammalian Total RNA kit (Sigma Aldrich) according to the manufacturer’s protocol (for more than 4×10^6^ cells) or Quick-RNA MicroPrep kit (Zymo research R1051, Irvine, California, USA) according to the manufacturer’s protocol (for fewer than 1×0^6^ cells). RNA was then transcribed to cDNA using the qScript cDNA Synthesis kit (Quantabio, Beverly, MA, USA) according to the manufacturer’s instructions, and RT-PCT or qRT-PCR was performed using the following primers:

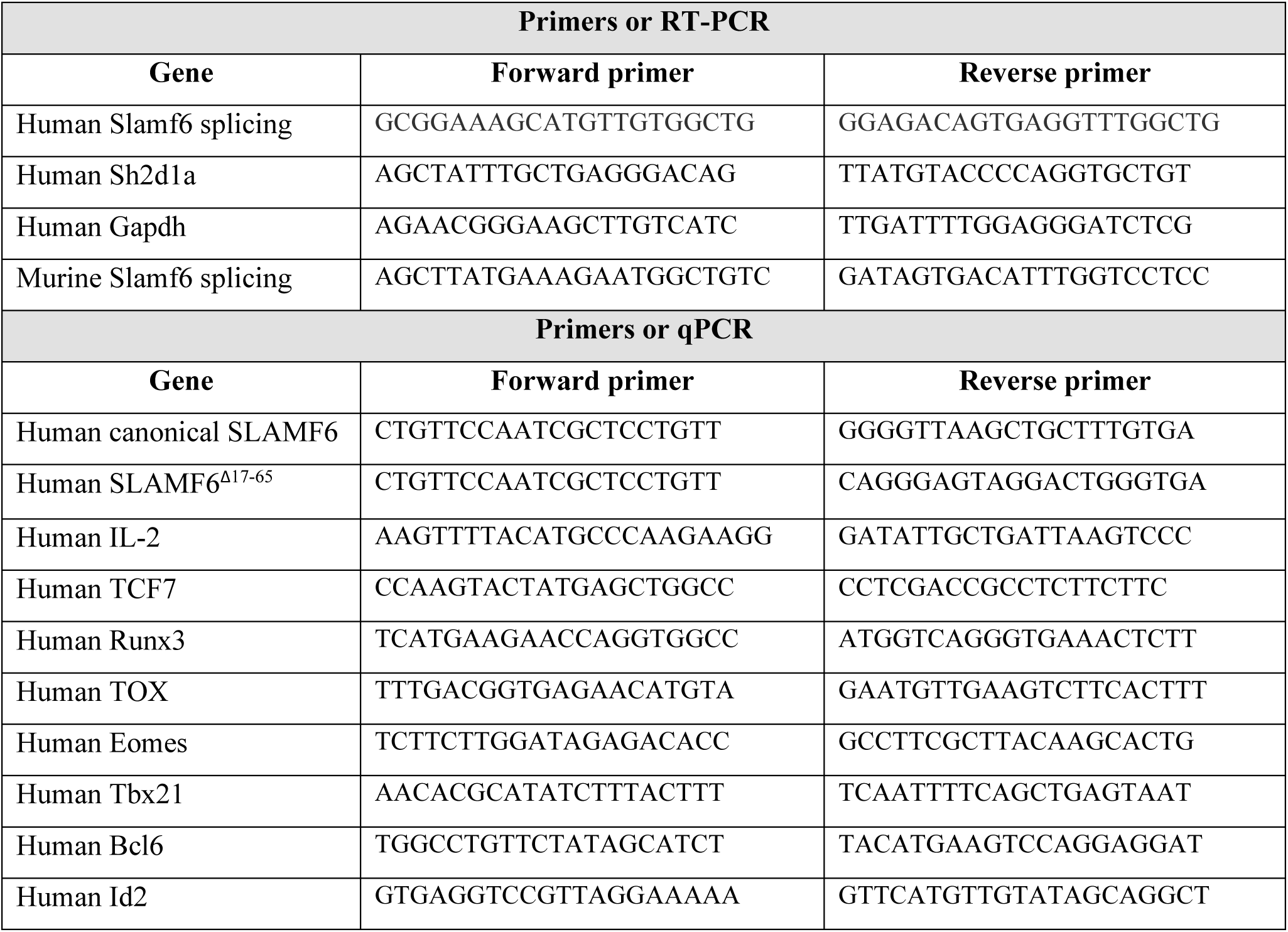
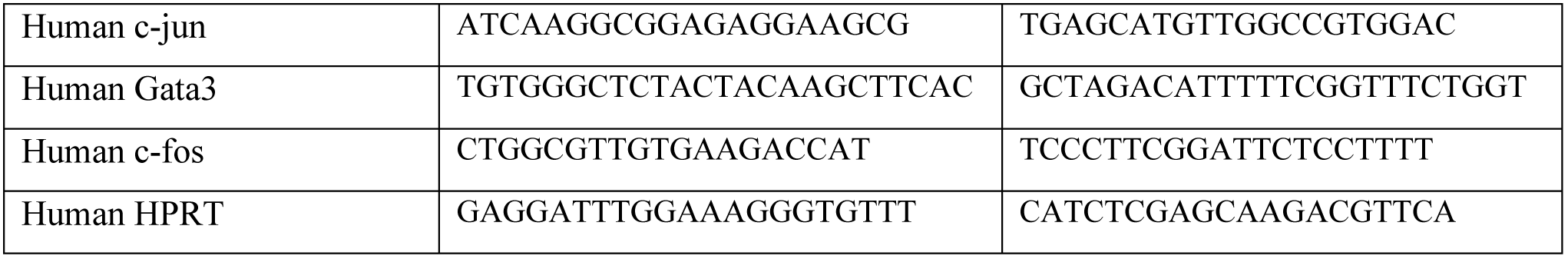

RT-PCR was performed in the SensQuest lab cycler machine (Danyel Biotech). The products were then run on 1.5% agarose gel, and qRT-PCR was performed in the CFX384 real-time system (Bio-Rad) using Bio-Rad CFX Manager 3.0 software.

#### Intracellular cytokine staining

TILs (1×10^5^) were co-cultured for 6h at 37°C at a 1:1 ratio with the indicated target melanoma cells. After 2h, brefeldin A (Biolegend) was added (1:1000). After incubation, the cells were washed twice with PBS and stained with anti-CD8 (Biolegend) for 30 minutes at room temperature. Following fixation and permeabilization (eBioscience protocol), intracellular IFN-γ and TNFα were labeled with anti-IFN-γ and anti-TNFα antibodies (Biologend), respectively for 30 minutes at room temperature. Cells were washed with permeabilization buffer, resuspended in FACS buffer, and subjected to flow cytometry.

#### Jurkat activation assay

1×10^5^ Jurkat cells (either WT or SLAMF6^Δ17-65^ cells) were activated with 200 ng/ml PMA and 300 ng/ml ionomycin for 48h. At the end of the activation, the conditioned medium was collected, and IL-2 or IFN-γ secretion was detected by ELISA (R&D).

#### Co-Culture experiment

Human lymphocytes (1×10^5^) were co-cultured overnight at a 1:1 ratio with the indicated target cells. Conditioned medium was collected, and IFN-γ secretion was detected by ELISA (R&D).

#### siRNA

siRNA against SLAMF6, SHP1 and SAP were purchased from QIAGEN (Hilden, Germany) (siSLAMF6(1) – SI00147252, siSLAMF6(2) – SI00147259, siSAP(1) – SI00036568, siSAP(2) – SI00036561, siSHP1(1) – SI02658726, siSHP1(2) – SI02658733). The cells were washed twice, resuspended in RPMI 1640 medium without supplements, and then electroporated in Biorad 0.2cm cuvettes using ECM 630 Electro Cell manipulator (BTX Harvard apparatus) at 250V, 300μF, 1000Ω. After electroporation, the cells were immediately seeded into complete RPMI medium. Twenty four hours post electroporation, the cells were activated and evaluated as described above.

#### Antisense oligos

ASOs were designed to target the splicing site in exon2; the oligos were purchased from Microsynth, Switzerland, synthetized with a full (all nucleotides) phosphorothioate (PS) backbone and in which each ribose 2′-hydroxyl was modified to 2′-methoxyethyl (2′-MOE). Transfection of Jurkat cells: Jurkat cells were collected, washed twice, resuspended in RPMI 1640 medium without supplements, and then electroporated in Biorad 0.2 cm cuvettes using ECM 630 Electro Cell manipulator (BTX Harvard apparatus) at 250V, 300μF, 1000Ω. After electroporation, the cells were immediately seeded into complete RPMI medium. Transfection of PBMCs: ASOs were electroporated into PBMCs using the Human T cell nucleofector kit (VPA-1002, Lonza, Basel, Switzerland). Transfection of TILs: 209TIL cells were electroporated according to the protocol^56^. All cells were evaluated twenty four hours post electroporation. Oligos sequences are as follows: Scrambled ASO: 5’ UGACCGAAAAGUCAUCUCAA 3’. ASO: 5’ GGGUACUAUGAAGGCAAGAG 3’.

#### Immunoblotting

Cells were lysed using RIPA buffer, and protein concentrations were measured using Bradford quantification. Equal concentrations of lysates were resuspended in SDS sample buffer (250 mM Tris-HCl [pH 6.8], 5% w/v SDS, 50% glycerol, and 0.06% w/v bromophenol blue) for 5 min at 95°C. Proteins were separated by SDS PAGE and transferred to a PVDF membrane. Membranes, blocked with 1% milk, were incubated with primary antibodies overnight at 4°C, followed by incubation with HRP-conjugated secondary antibodies for 1 h at RT (Jackson ImmunoResearch Laboratories). Signals were detected by enhanced chemiluminescence reagents (Clarity Western ECL Substrate, BioRad, Hercules, CA, USA).

#### Binding competition assay

Cells were incubated with increasing concentrations from 25 to 100 µg/ml seSLAMF6 for 30 min on ice, washed twice and labeled with Goat anti SLAMF6 (R&D), and a secondary Ab (Donkey anti Goat, Jackson ImmunoResearch) on ice. Antibody binding was measured by flow cytometry.

#### Biotinylation of soluble ectodomain proteins

seSLAMF6 and seSLAMF6^Δ17-65^ were biotinylated using EZ Link Sulfo NHS Biotin (ThermoFisher Scientific) for 30 min at room temperature. Biotinylated proteins were collected using 15 min centrifugation at 1700rcf in an Amicon® Ultra-4 Centrifugal Filter Unit (Millipore) in 2ml PBS.

#### ELISA for binding capacity of splice isoforms

A plate was coated overnight with 0.5μM soluble SLAMF6 ectodomain or soluble SLAMF6^Δ17-65^ ectodomain in coating buffer (8.4g NaHCO_3_, 3.56g Na_2_CO_3_ in 1L DDW, pH 9.5), 4°C. The next day the plate was washed and blocked with blocking buffer (1%BSA in PBS) for 1 h at room temperature and then biotinylated proteins (seSLAMF6 or seSLAMF6^Δ17-65^) were added for 2 h at room temperature. Streptavidin-HRP was added for 1 h at room temperature. Signals were detected by the addition of TMB/E substrate (ES001, Millipore).

#### Flow cytometry

Cells were stained with antibodies at room temperature for 25 min, according to the manufacturer’s instructions. Subsequently, cells were washed and analyzed using CytoFlex (Beckman Coulter, CA, USA), and flow cytometry-based sorting was performed in an ARIA-III sorter. Flow cytometry analysis was performed using FCS Express 5 flow research edition (De Novo software).

#### Statistics

Statistical significance was determined by unpaired t-test (two-tailed with equal SD) using Prism software (GraphPad). A p-value of < 0.05 was considered statistically significant. Analysis of more than twogroups was performed using a one-way ANOVA test. *, p≤0.05; **, p≤0.01; ***, p≤0.001. For each experiment, the number of replicates and the statistical test used are stated in the corresponding figure legend.

### RNA sequencing

#### RNA sequencing analysis for public datasets

Datasets with the following accession numbers were downloaded from NCBI SRA: SRR7588547, SRR7588548, SRR7588549, SRR7588550, SRR7588559, SRR7588560, SRR7588561, SRR7588562, SRR7588567, SRR7588568, SRR7588573, SRR7588574, SRR7588579, SRR7588580.

#### Bioinformatics

Poly-A/T stretches and Illumina adapters were trimmed from the reads using cutadapt^57^; resulting reads shorter than 30bp were discarded. Reads were mapped to the *H. sapiens* reference genome hg38 using STAR^58^, supplied with gene annotations downloaded from RefSeq^59^ (and with EndToEnd option and outFilterMismatchNoverLmax set to 0.04). Isoform expression was analyzed with the RSEM version 1.3.0^60^. The pipeline was run using snakemake^61^. The genome browser was Jbrowse (https://www.ncbi.nlm.nih.gov/pmc/articles/PMC4830012/).

#### RNA sequencing of activated WT and SLAMF6^Δ17-65^ Jurkat cells

WT and SLAMF6^Δ17-65^ Jurkat cells activated (0, 6h, 12h, 24h and 36h) in 96 flat bottom plates in the presence or absence of PMA (200ng/ml) and ionomycin (300ng/ml) were lysed in 100Sµl lysis buffer containing TCL buffer (Qiagen, 1031576) containing 1% β-mercaptoethanol. After lysis, plates were imminently frozen and stored at -80°C until samples were processed. Libraries were created using the Smart-Seq2 protocol as previously described^62^, with minor modifications for bulk populations. For RNA purification 10µl of cell lysates for each sample was used. 40µl of Agencourt RNAClean XP SPRI beads (Beckman Coulter, A63987) were added to the cell lysates, and after 10 minutes of incubation at room temperature, plates were placed on a DynaMag 96 side skirted magnet (Invitrogen, 12027) for 5 minutes, followed by supernatant removal and 2 cycles of 75% ethanol washes. After ethanol removal, plates were incubated for 15 minutes or until beads started to crack, and 4µl of Mix-I (containing: 1µl (10µM) RT primer (DNA oligo) 5′–AAGCAGTGGTATCAACGCAGAGTACT30VN-3′; 1µl (10mM) dNTPs; 1µl (10%, 4U/µl) Recombinant RNase Inhibitor; and 1µl nuclease-free water) was added and mixed in each well, following incubation at 72°C for 3 minutes. Next, 7µl of Mix-II (containing: 0.75µl nuclease free water; 2µl 5X Maxima RT buffer; 2µl (5M) betaine; 0.9µl (100mM) MgCl2; 1µl (10µM) TSO primer (RNA oligo) 5′-AAGCAGTGGTATCAACGCAGAGTACATrGrG+G-3′; 0.25µl (40U/µl) Recombinant RNase Inhibitor; and 0.1µl (200U/µl) of Maxima H Minus Reverse Transcriptase), was added and mixed, and reverse transcription (RT) was performed at 50°C for 90 minutes, followed by 5 minutes incubation at 85°C. After RT incubation, 14µl of Mix-III (containing: 1µl nuclease free water; 0.5µl (10µM) ISPCR primer (DNA oligo) 5′-AAGCAGTGGTATCAACGCAGAGT-3′; and 12.5µl 2X KAPA HiFi HotStart ReadyMix), was added for the cDNA amplification step, performed at 98°C for 3 minutes, followed by 12 cycles at [98°C for 15 seconds, 67°C for 20 seconds and 72°C for 6 minutes] with a final extension at 72°C for 5 minutes. Next, two cycles for removal of primer dimer residues were performed by adding 20µl of Agencourt AMPureXP SPRI beads (Beckman Coulter, A63881), incubation of plates for 5 minutes at room temperature, followed by 5 minutes of incubation on a DynaMag 96 side skirted magnet, 2 cycles of 75% ethanol washes, and resuspension of dried beads with 50µl of TE buffer (Tekanova, T0228). After generating the cDNA, quality control was performed in order to evaluate 1) the concentration of each sample using the Qubit dsDNA high sensitivity assay kit (Life Technologies, Q32854), and 2) size distribution using the High Sensitivity DNA BioAnalyzer kit (Agilent, 5067-4626). Libraries were constructed using the Nextera XT library Prep kit (Illumina, FC-131-1096), and combined libraries were sequenced on a NextSeq 500 sequencer (Illumina) using the 75 cycles kit, with paired-end 38-base-reads and dual barcoding.

#### Data analysis

FASTQ files were aligned to the NCBI Human Reference Genome Build GRCh37 (hg19) using STAR^58^. Principal component analysis (PCA) was performed using the DeSeq2 plot PCA function. Differential expression was performed using DeSeq2 results function, with lfcThreshold = 0.25 and altHypothesis = “greaterAbs”. The Benjamini-Hochberg procedure corrected p-values. Significant genes were chosen if they had adjusted p-values <0.05. Differential expression was calculated either comparing each timepoint to its corresponding sample type at 0 h or as differentially expressed genes between WT and Δ17-65 samples for each time point. Expression values were normalized via the DESeq2 DESeq function. Next, each value was subtracted by the mean of its gene expression in all the samples. Immune-related genes were selected from the RNA sequencing results based on a unified list of genes created from the Immunogenetic Related Information Source (IRIS) list and the MAPK/NFKB Network list (https://www.innatedb.com/redirect.do?go=resourcesGeneLists). Sashimi plots were generated with the IGV visualization tool^63^.

#### Enrichment analysis

Genes that were highly expressed compared to 0 h for each cell type were analyzed by GeneAnalytics. The scores were calculated as the -log2(p-value). (http://geneanalytics.genecards.org)^64^.

#### Cytotoxic score

The cytotoxic score was calculated by using a method similar to that of Tirosh et al.^25^ with the same naϊve and cytotoxic genes chosen. The value for each gene was calculated as the average expression of three replicates of a sample type at a specific time point.

## *In vivo* assay

### *Winn assay*. 209*TIL*

cells electroporated with scrambled ASO or ASO1 were washed and mixed at a 1:1 ratio (1×10^6^ cells each) with 526*mel* and immediately injected subcutaneously into the backs of 8- to 9-week-old female nude (athymic Foxn1^-/-^) mice (N=7). Tumor size was measured in two perpendicular diameters three times per week. Mice were sacrificed when tumors reached a 15 mm diameter in one dimension or when the lesion necrotized. Tumor volume was calculated as L(length) x W (width)^2^ x 0.5.

Animal studies were approved by the Institutional Review Board - Authority for biological and biomedical models, Hebrew University, Jerusalem, Israel. (MD-20-16104-5).

## Data availability

The RNAseq data generated in this work is being uploaded to data repositories (we will update the statement as soon as we have the accession numbers).

The processed gene expression data is deposited to GEO, the raw sequencing files are deposited to dbGap.

## Acknowledgments

We wish to acknowledge the devoted technical work of Inna Ben-David, Anna Kuznetz, and Yael Gelfand. We thank Eli Pikarsky and Ofer Mandelboim for helpful discussions, and Karen Pepper for editing the manuscript. We thank the Mantoux Institute for Bioinformatics of the Nancy and the Stephen Grand Israel National Center for Personalized Medicine, Weizmann Institute of Science, for data analysis. We are very grateful to Stewart and Maggie Greenberg for their long-standing support.

## Authors contributions

E.H. contributed to the conception, design of the work, acquisition, analysis and interpretation of the data and drafted the work. E.Z. contributed to the analysis and interpretation of the data. S.T. contributed to the acquisition and alaysis of the data. S.M. contributed to the analysis and interpretation of the data. J.C. contributed to the analysis and interpretation of the data. S.K. contributed to the analysis and interpretation of the data. S.F. contributed to the analysis and interpretation of the data. M.S.F contributed to the acquisition and alaysis of the data. Y.T. contributed to the analysis and interpretation of the data. K.Y. contributed to the analysis and interpretation of the data. A.M. contributed to the interpretation of the data. P.S. contributed to the acquisition of the data. N.H. contributed to the analysis and interpretation of the data. T.P. contributed to the analysis and interpretation of the data. A.R. contributed to the conception and design of the data and substantially revised the work. R.K. contributed to the conception and design of the data and substantially revised the work. G.E. contributed to the conception and design of the data and substantially revised the work. M.L. contributed to the conception and design of the data and substantially revised the work.

## References

1. Wu, N. & Veillette, A. SLAM family receptors in normal immunity and immune pathologies. Curr Opin Immunol 38, 45–51, doi: 10.1016/j.coi.2015.11.003 (2016).

2. Cannons, J. L., Tangye, S. G. & Schwartzberg, P. L. SLAM family receptors and SAP adaptors in immunity. Annu Rev Immunol 29, 665–705, doi: 10.1146/annurev-immunol-030409-101302 (2011).

3. Zhao, F., Cannons, J. L., Dutta, M., Griffiths, G. M. & Schwartzberg, P. L. Positive and negative signaling through SLAM receptors regulate synapse organization and thresholds of cytolysis. Immunity 36, 1003–1016, doi: 10.1016/j.immuni.2012.05.017 (2012).

4. Biram, A., Davidzohn, N. & Shulman, Z. T cell interactions with B cells during germinal center formation, a three-step model. Immunol Rev 288, 37–48, doi: 10.1111/imr.12737 (2019).

5. Li, W. et al. The SLAM-associated protein signaling pathway is required for development of CD4+ T cells selected by homotypic thymocyte interaction. Immunity 27, 763–774, doi: 10.1016/j.immuni.2007.10.008 (2007).

6. Guo, H. et al. Deletion of Slam locus in mice reveals inhibitory role of SLAM family in NK cell responses regulated by cytokines and LFA-1. J Exp Med 213, 2187–2207, doi: 10.1084/jem.20160552 (2016).

7. Lu, Y. et al. SLAM receptors foster iNKT cell development by reducing TCR signal strength after positive selection. Nat Immunol 20, 447–457, doi: 10.1038/s41590-019-0334-0 (2019).

8. Shlapatska, L. M. et al. CD150 association with either the SH2-containing inositol phosphatase or the SH2-containing protein tyrosine phosphatase is regulated by the adaptor protein SH2D1A. J Immunol 166, 5480–5487, doi: 10.4049/jimmunol.166.9.5480 (2001).

9. Morra, M. et al. Structural basis for the interaction of the free SH2 domain EAT-2 with SLAM receptors in hematopoietic cells. EMBO J 20, 5840–5852, doi: 10.1093/emboj/20.21.5840 (2001).

10. Latour, S. et al. Regulation of SLAM-mediated signal transduction by SAP, the X-linked lymphoproliferative gene product. Nat Immunol 2, 681–690, doi: 10.1038/90615 (2001).

11. Eissmann, P. et al. Molecular basis for positive and negative signaling by the natural killer cell receptor 2B4 (CD244). Blood 105, 4722–4729, doi: 10.1182/blood-2004-09-3796 (2005).

12. Snow, A. L. et al. Restimulation-induced apoptosis of T cells is impaired in patients with X-linked lymphoproliferative disease caused by SAP deficiency. J Clin Invest 119, 2976–2989, doi: 10.1172/JCI39518 (2009).

13. Hajaj, E. et al. SLAMF6 deficiency augments tumor killing and skews toward an effector phenotype revealing it as a novel T cell checkpoint. Elife 9, doi: 10.7554/eLife.52539 (2020).

14. Zhong, M. C. & Veillette, A. Control of T lymphocyte signaling by Ly108, a signaling lymphocytic activation molecule family receptor implicated in autoimmunity. J Biol Chem 283, 19255–19264, doi: 10.1074/jbc.M800209200 (2008).

15. Kumar, K. R. et al. Regulation of B cell tolerance by the lupus susceptibility gene Ly108. Science 312, 1665–1669, doi: 10.1126/science.1125893 (2006).

16. Bottino, C. et al. NTB-A [correction of GNTB-A], a novel SH2D1A-associated surface molecule contributing to the inability of natural killer cells to kill Epstein-Barr virus-infected B cells in X-linked lymphoproliferative disease. J Exp Med 194, 235–246, doi: 10.1084/jem.194.3.235 (2001).

17. Fraser, C. C. et al. Identification and characterization of SF2000 and SF2001, two new members of the immune receptor SLAM/CD2 family. Immunogenetics 53, 843–850, doi: 10.1007/s00251-001-0415-7 (2002).

18. O’Leary, N. A. et al. Reference sequence (RefSeq) database at NCBI: current status, taxonomic expansion, and functional annotation. Nucleic Acids Res 44, D733–745, doi: 10.1093/nar/gkv1189 (2016).

19. Cichocki, F. et al. ARID5B regulates metabolic programming in human adaptive NK cells. J Exp Med 215, 2379–2395, doi: 10.1084/jem.20172168 (2018).

20. Calpe, S. et al. The SLAM and SAP gene families control innate and adaptive immune responses. Adv. Immunol. 97, 177–250 (2008)

21. Veillette, A. SLAM-family receptors: immune regulators with or without SAP-family adaptors. Cold Spring Harb. Perspect. Biol. 2, a002469 (2010)

22. Yigit, B. et al. SLAMF6 as a Regulator of Exhausted CD8 T Cells in Cancer. Cancer Immunology Research vol. 7 1485–1496 (2019)

23. Eisenberg, G. et al. Soluble SLAMF6 Receptor Induces Strong CD8 T-cell Effector Function and Improves Anti-Melanoma Activity In Vivo. Cancer Immunology Research vol. 6 127–138 (2018)

24. Schweingruber, C., Rufener, S. C., Zünd, D., Yamashita, A. & Mühlemann, O. Nonsense-mediated mRNA decay — Mechanisms of substrate mRNA recognition and degradation in mammalian cells. Biochimica et Biophysica Acta (BBA) - Gene Regulatory Mechanisms vol. 1829 612–623 (2013)

25. Tirosh, I. et al. Dissecting the multicellular ecosystem of metastatic melanoma by single-cell RNA-seq. Science 352, 189–196 (2016)

26. Dupre, L. et al. SAP controls the cytolytic activity of CD8+ T cells against EBV-infected cells. Blood 105, 4383–4389, doi: 10.1182/blood-2004-08-3269 (2005).

27. Latour, S. et al. Binding of SAP SH2 domain to FynT SH3 domain reveals a novel mechanism of receptor signalling in immune regulation. Nat Cell Biol 5, 149–154, doi: 10.1038/ncb919 (2003).

28. Panchal, N., Booth, C., Cannons, J. L. & Schwartzberg, P. L. X-Linked Lymphoproliferative Disease Type 1: A Clinical and Molecular Perspective. Front Immunol 9, 666, doi: 10.3389/fimmu.2018.00666 (2018).

29. Kageyama, R. et al. The receptor Ly108 functions as a SAP adaptor-dependent on-off switch for T cell help to B cells and NKT cell development. Immunity 36, 986–1002, doi: 10.1016/j.immuni.2012.05.016 (2012).

30. Hodi, F. S. et al. Improved survival with ipilimumab in patients with metastatic melanoma. N. Engl. J. Med. 363, 711–723 (2010)

31. Topalian, S. L. et al. Survival, durable tumor remission, and long-term safety in patients with advanced melanoma receiving nivolumab. J. Clin. Oncol. 32, 1020–1030 (2014)

32. Wheeler, T. M. et al. Targeting nuclear RNA for in vivo correction of myotonic dystrophy. Nature 488, 111–115 (2012)

33. Porensky, P. N. & Burghes, A. H. Antisense oligonucleotides for the treatment of spinal muscular atrophy. Hum Gene Ther 24, 489–498, doi: 10.1089/hum.2012.225 (2013).

34. Kozlovski, I., Siegfried, Z., Amar-Schwartz, A. & Karni, R. The role of RNA alternative splicing in regulating cancer metabolism. Hum Genet 136, 1113–1127, doi: 10.1007/s00439-017-1803-x (2017).

35. Oltean, S. & Bates, D. O. Hallmarks of alternative splicing in cancer. Oncogene 33, 5311–5318, doi: 10.1038/onc.2013.533 (2014).

36. Ergun, A. et al. Differential splicing across immune system lineages. Proc Natl Acad Sci U S A 110, 14324–14329, doi: 10.1073/pnas.1311839110 (2013).

37. Rotival, M., Quach, H. & Quintana-Murci, L. Defining the genetic and evolutionary architecture of alternative splicing in response to infection. Nat Commun 10, 1671, doi: 10.1038/s41467-019-09689-7 (2019).

38. West, K. O. et al. The Splicing Factor hnRNP M Is a Critical Regulator of Innate Immune Gene Expression in Macrophages. Cell Rep 29, 1594–1609 e1595, doi: 10.1016/j.celrep.2019.09.078 (2019).

39. Ye, C. J. et al. Genetic analysis of isoform usage in the human anti-viral response reveals influenza-specific regulation of ERAP2 transcripts under balancing selection. Genome Res 28, 1812–1825, doi: 10.1101/gr.240390.118 (2018).

40. Shemesh, A. et al. First Trimester Pregnancy Loss and the Expression of Alternatively Spliced NKp30 Isoforms in Maternal Blood and Placental Tissue. Front Immunol 6, 189, doi: 10.3389/fimmu.2015.00189 (2015).

41. Oberdoerffer, S. et al. Regulation of CD45 alternative splicing by heterogeneous ribonucleoprotein, hnRNPLL. Science 321, 686–691 (2008)

42. LaFleur, M. W. et al. PTPN2 regulates the generation of exhausted CD8(+) T cell subpopulations and restrains tumor immunity. Nat Immunol 20, 1335–1347, doi: 10.1038/s41590-019-0480-4 (2019).

43. Miller, B. C. et al. Subsets of exhausted CD8(+) T cells differentially mediate tumor control and respond to checkpoint blockade. Nat Immunol 20, 326–336, doi: 10.1038/s41590-019-0312-6 (2019).

44. Chen, Z. et al. TCF-1-Centered Transcriptional Network Drives an Effector versus Exhausted CD8 T Cell-Fate Decision. Immunity 51, 840–855 e845, doi: 10.1016/j.immuni.2019.09.013 (2019).

45. Wei, J. et al. Targeting REGNASE-1 programs long-lived effector T cells for cancer therapy. Nature 576, 471–476, doi: 10.1038/s41586-019-1821-z (2019).

46. Martinez-Usatorre, A. et al. Enhanced Phenotype Definition for Precision Isolation of Precursor Exhausted Tumor-Infiltrating CD8 T Cells. Front Immunol 11, 340, doi: 10.3389/fimmu.2020.00340 (2020).

47. Chemnitz, J. M., Parry, R. V., Nichols, K. E., June, C. H. & Riley, J. L. SHP-1 and SHP-2 associate with immunoreceptor tyrosine-based switch motif of programmed death 1 upon primary human T cell stimulation, but only receptor ligation prevents T cell activation. J Immunol 173, 945–954, doi: 10.4049/jimmunol.173.2.945 (2004).

48. Peled, M. et al. Affinity purification mass spectrometry analysis of PD-1 uncovers SAP as a new checkpoint inhibitor. Proc Natl Acad Sci U S A 115, E468–E477, doi: 10.1073/pnas.1710437115 (2018).

49. Meazza, R. et al. XLP1 inhibitory effect by 2B4 does not affect DNAM-1 and NKG2D activating pathways in NK cells. Eur J Immunol 44, 1526–1534, doi: 10.1002/eji.201344312 (2014).

50. Wu, N. et al. A hematopoietic cell–driven mechanism involving SLAMF6 receptor, SAP adaptors and SHP-1 phosphatase regulates NK cell education. Nat. Immunol. 17, 387–396 (2016)

51. Clemens, P. R. et al. Safety, Tolerability, and Efficacy of Viltolarsen in Boys With Duchenne Muscular Dystrophy Amenable to Exon 53 Skipping: A Phase 2 Randomized Clinical Trial. JAMA Neurol, doi: 10.1001/jamaneurol.2020.1264 (2020).

52. Mercuri, E. et al. Nusinersen versus Sham Control in Later-Onset Spinal Muscular Atrophy. N Engl J Med 378, 625–635, doi: 10.1056/NEJMoa1710504 (2018).

53. Havens, M. A. & Hastings, M. L. Splice-switching antisense oligonucleotides as therapeutic drugs. Nucleic Acids Res 44, 6549–6563, doi: 10.1093/nar/gkw533 (2016).

54. Lundin, K. E., Gissberg, O., Smith, C. I. E. & Zain, R. Chemical Development of Therapeutic Oligonucleotides. Methods Mol Biol 2036, 3–16, doi: 10.1007/978-1-4939-9670-4_1 (2019).

55. Ran, F. A. et al. Genome engineering using the CRISPR-Cas9 system. Nat. Protoc. 8, 2281–2308 (2013)

56. Transfection of mammalian cells by electroporation. Nat. Methods 3, 67–68 (2006)

57. Martin, M. Cutadapt removes adapter sequences from high-throughput sequencing reads. EMBnet.journal vol. 17 10 (2011)

58. Dobin, A. et al. STAR: ultrafast universal RNA-seq aligner. Bioinformatics 29, 15–21 (2013)

59. Brister, J. R., Ako-Adjei, D., Bao, Y. & Blinkova, O. NCBI viral genomes resource. Nucleic Acids Res. 43, D571–7 (2015)

60. Köster, J. & Rahmann, S. Snakemake—a scalable bioinformatics workflow engine. Bioinformatics vol. 34 3600–3600 (2018)

61. Li, B. & Dewey, C. N. RSEM: accurate transcript quantification from RNA-Seq data with or without a reference genome. BMC Bioinformatics vol. 12 (2011)

62. Sade-Feldman, M. et al. Defining T Cell States Associated with Response to Checkpoint Immunotherapy in Melanoma. Cell 176, 404 (2019)

63. Fuchs, S. B.-A. et al. GeneAnalytics: An Integrative Gene Set Analysis Tool for Next Generation Sequencing, RNAseq and Microarray Data. OMICS: A Journal of Integrative Biology vol. 20 139–151 (2016)

64. James T. Robinson, Helga Thorvaldsdóttir, Wendy Winckler, Mitchell Guttman, Eric S. Lander, Gad Getz, Jill P. Mesirov. Integrative Genomics Viewer. Nature Biotechnology 29, 24–26 (2011)

