## Supplementary figures for "Alternative splicing of SLAMF6 in human T cells creates a co-stimulatory isoform that counteracts the inhibitory effect of the full-length receptor"

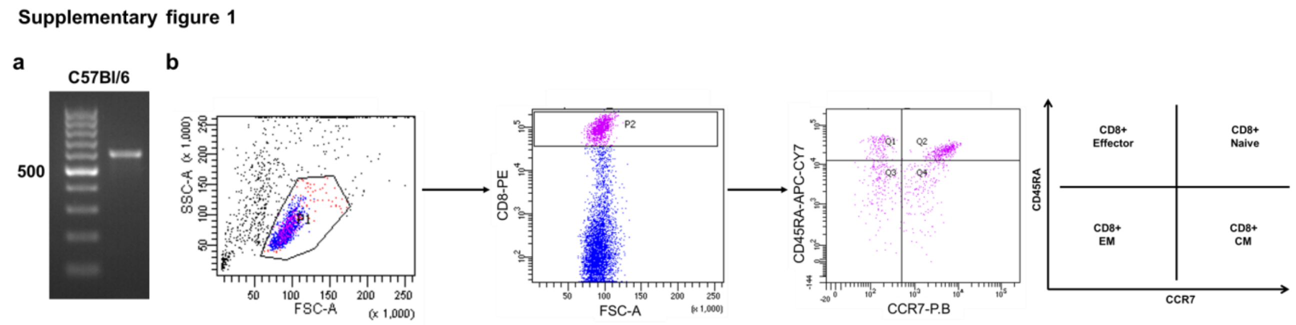


**Supplementary figure 1:**

**a**, RNA expression of *LY108*, the murine analog of *SLAMF6*, in C57Bl/6 splenocytes. **b**, Gating strategy for cell sorting separating CD8 T cells to subsets according to CD45RA/CCR7 expression.

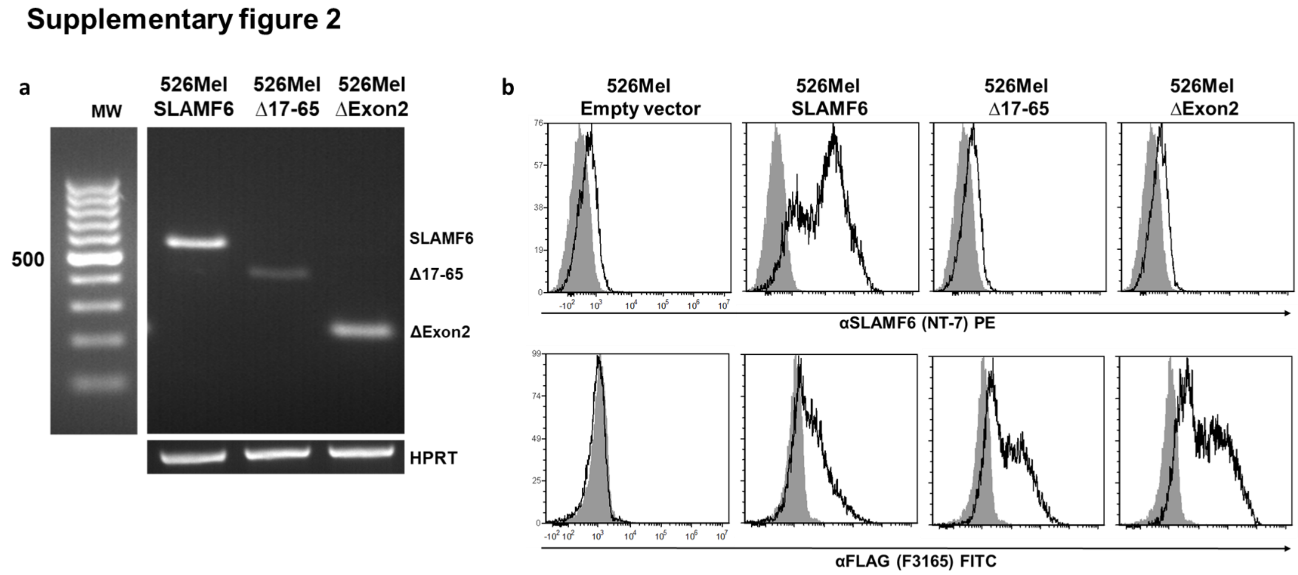


**Supplementary figure 2:**

Validation of SLAMF6 protein expression in modified melanoma cells. **a**, RNA expression of *SLAMF6* isoforms on transfected 526*mel* melanoma cells. **b**, Flow cytometry to determine SLAMF6 (MoAb NT-7) or FLAG (MoAb F3165) expression on the transfected 526*mel* cells.

Supplementary figure 3

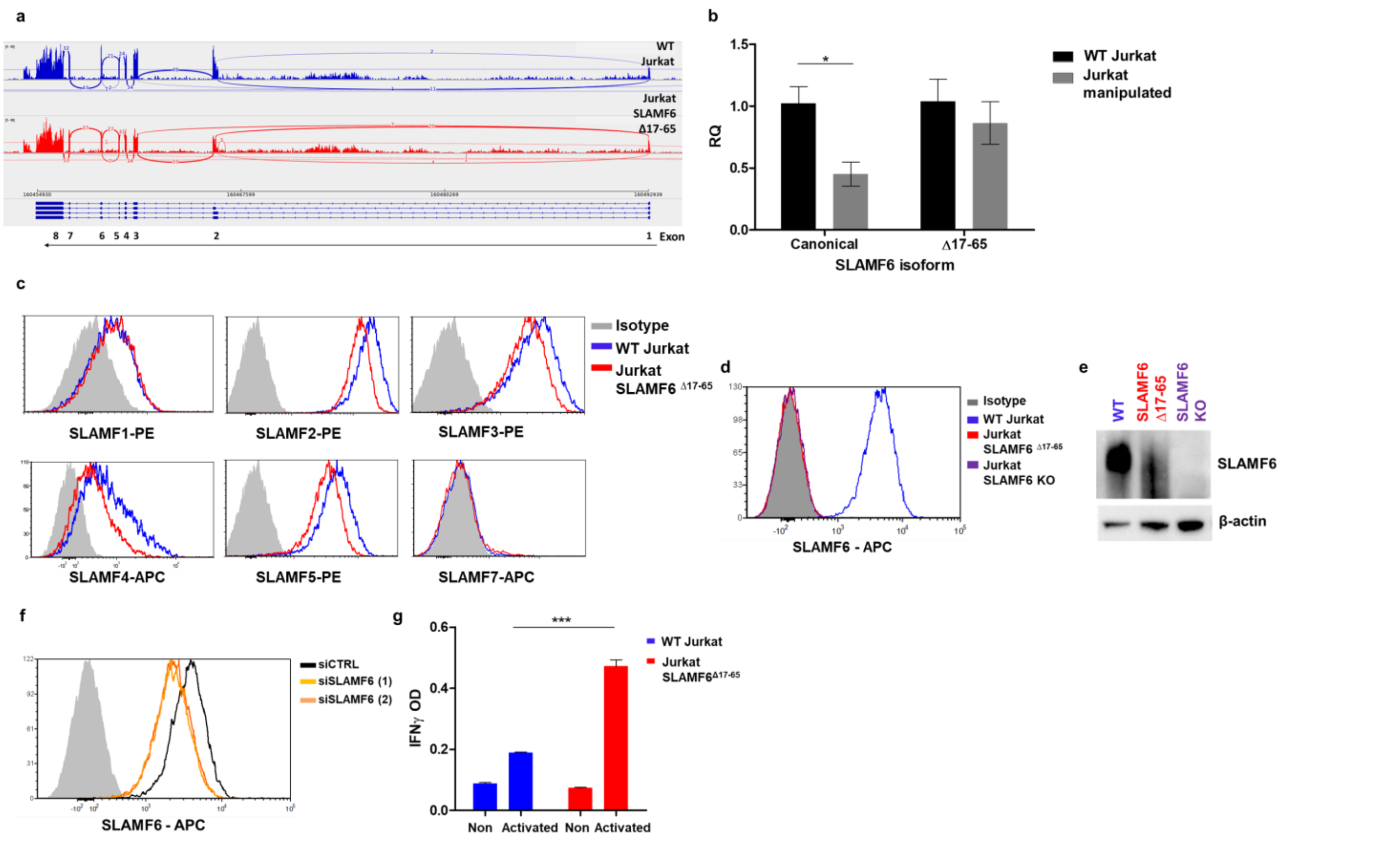


**Supplementary figure 3:**

**a**, Sashimi plot showing SLAMF6 splice isoforms in WT Jurkat cells and Jurkat-$SLAMF6^{\Delta17\text{-}65}$ cells 36 h post-activation. **b**, Quantitative RT-PCR for *SLAMF6-canonical* and *$SLAMF6^{\Delta17\text{-}65}$* in relevant Jurkat cells. Data were normalized to *HPRT* expression and WT Jurkat expression. **c**, Expression of SLAM family members in WT Jurkat and Jurkat-$SLAMF6^{\Delta17\text{-}65}$ cells. **d**, SLAMF6 canonical isoform expression in three Jurkat cell lines: WT, $SLAMF6^{\Delta17\text{-}65}$ , and SLAMF6 KO. **e**, Immunoblot of SLAMF6 using anti-SLAMF6 (AF1908) targeting SLAMF6 C-domain. **f**, SLAMF6 canonical isoform silencing in WT Jurkat cells electroporated with siSLAMF6. **g**, ELISA for IFN-γ in WT Jurkat and Jurkat-$SLAMF6^{\Delta17\text{-}65}$ cells activated for 48 h. OD, optical density. One-way ANOVA. ***, $P < 0.001$.

Supplementary figure 4

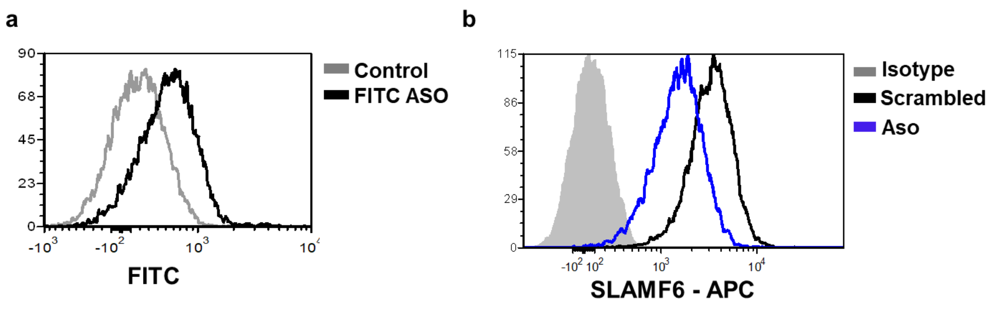


**Supplementary figure 4:**

**a**, FITC levels measured by flow cytometry 24 h post electroporation of WT Jurkat cells with ASO-FITC or control ASO. **b**, SLAMF6 expression measured using flow cytometry 24 h post-electroporation of WT Jurkat cells with ASO or control ASO.
